# Multi-strain SIS dynamics with coinfection across heterogeneous patches

**DOI:** 10.1101/2025.10.26.684704

**Authors:** Miguel B. Marôco, Louis Pailloncy, Sten Madec, Erida Gjini

## Abstract

We investigate a spatially structured Susceptible–Infected–Susceptible (SIS) model with coinfection, multiple interacting strains, and host movement across a network of heterogeneous patches. Motivated by the need to capture how local epidemiological processes and spatial connectivity jointly shape multi-strain pathogen dynamics, we consider a metacommunity framework in which hosts migrate between interconnected habitats. Under assumptions of strain similarity and slow migration, and following the continuous-space formulation of Le and Madec (2023), we derive a reduced discrete model that separates fast epidemiological dynamics from slow strain dynamics. The fast variables describe the total densities of susceptible, singly infected, and co-infected hosts within each patch, while the slow variables track strain frequencies locally. These frequencies evolve according to a replicator-like equation that combines local selection with an explicit migration term modulated by patch heterogeneity. The model allows *N* strains to differ across multiple epidemiological traits-such as transmission, clearance, and coinfection interactions - both within and between *P* patches, leading to complex and potentially non-intuitive system behavior. By leveraging the analytical tractability of the reduced framework, we characterize how spatial structure, host movement, and environmental heterogeneity influence key ecological outcomes, including coexistence, competitive exclusion, and spatial segregation of strains. Illustrating numerically low-dimensional cases (*N* = 2, *P* = 2, 3), we show how the interplay between local dynamics and spatial coupling shapes global system behavior. More broadly, this framework provides a tractable and mechanistically grounded approach for studying multi-strain pathogen dynamics in heterogeneous environments, contributing to ongoing efforts to bridge ecological realism, computational feasibility, and analytical insight in spatial ecology and infectious diseases.

## 1. Introduction

Population dynamics in heterogeneous environments are central to biology, ecology, epidemiology, and medicine, with applications ranging from ecosystem stability and metapopulation persistence to infectious disease spread and biodiversity conservation (Hanski, 1999; Okubo et al., 2001; Keeling et al., 2004; Hanski and Ovaskainen, 2000; Tao et al., 2024). Classical modeling frameworks—such as logistic growth with diffusion, Lotka–Volterra systems with migration, and epidemiological compartmental models across connected environments—have long been used to study how populations evolve across space (Arditi et al., 2018; Levin, 1974; Smith et al., 2004). Similar questions arise in within-host systems, where pathogens spread across tissues or organs (Wong and Yukl, 2016; Grant et al., 2008). Across these contexts, key theoretical questions concern persistence, stability, and how diffusion and environmental heterogeneity or landscape fragmentation shape system dynamics and processes (Butler et al., 1986; Arditi et al., 2015; Poggiale et al., 2005; Hanski and Ovaskainen, 2000; Tao et al., 2024).

In epidemiology, spatial disease modeling has evolved from early reaction–diffusion approaches to more realistic multi-patch frameworks that represent discrete populations connected by movement (Noble, 1974; Capasso and Maddalena, 1981; Murray et al., 1986; McCormack and Allen, 2007). Even simple spatial extensions can fundamentally alter disease persistence and spread (Skellam, 1951), while heterogeneity in space, mobility, and transmission processes greatly increases system complexity. These developments have contributed to the emergence of spatial infectious disease ecology as a mature and interdisciplinary field (Pullan et al., 2012; Parratt et al., 2016; Lin and Wen, 2022).

Despite major advances in high-resolution data, computational tools, and modeling approaches, significant challenges remain. A central issue is the mismatch between model requirements and available data: surveillance systems are often incomplete or biased (Kraemer et al., 2016; Lin and Wen, 2022). At the same time, disease dynamics operate across multiple spatial and temporal scales, from within-host processes to global transmission networks, making it difficult to develop models that consistently bridge these scales (Meentemeyer et al., 2012; Real and Biek, 2007; Malagon et al., 2025; Zilio et al., 2026). Rapid environmental change, driven by climate and land-use shifts, further complicates prediction by violating assumptions of stationarity that underpin many classical models.

Another major challenge lies in capturing the complexity of host–pathogen-environment interactions. Disease dynamics are shaped by nonlinear feedbacks involving population structure, human mobility, and pathogen evolution, including phylodynamics. While recent advances have generated large and diverse datasets-from remote sensing, genomics, and mobility tracking - integrating these heterogeneous sources remains technically demanding due to differences in scale, uncertainty, and bias. For instance, linking pathogen phylogenies with spatial transmission models is still an active area of research. More broadly, there is an inherent tension between model realism and tractability: models must be sufficiently detailed to capture complex spatial dynamics, yet simple and interpretable enough to support inference, prediction, and decision-making.

Motivated by these challenges, where pathogen diversity and host and environmental heterogeneity intertwine, spatially structured (multi-patch) multi-strain models provide a natural framework for studying disease dynamics in realistic settings. Many populations-human, livestock, and wildlife-are organized into interconnected subpopulations, or “patches,” linked by movement. Increasing global connectivity has amplified the importance of these structures, enabling pathogens and their variants to spread rapidly across regions and scales (Steiner and Novembre, 2022; Plümper and Neumayer, 2023; Belik et al., 2011). Capturing the interplay between local transmission dynamics, host movement, and environmental heterogeneity is therefore essential for understanding modern disease ecology.

In this work, we build on recent advances in multi-strain epidemiological modeling (Madec and Gjini, 2020; Le et al., 2023; Le and Madec, 2023) to study a multi-patch, multi-strain SIS system with coinfection and host migration. In non-spatial settings, such systems can be reduced to replicator equations governing strain frequencies, capturing slow evolutionary dynamics driven by pairwise invasion fitnesses. Extending this framework to discrete heterogeneous patches allows us to incorporate both local selection and movement-driven effects in a unified and tractable way.

Our aims are threefold: (i) to derive and present the model reduction governing equations for strain frequencies across patches, (ii) to analyze key low-dimensional cases highlighting how patch heterogeneity and strain variability interact to shape selection, and (iii) to provide a foundation for further theoretical and applied extensions. By combining ecological realism with analytical tractability, this approach contributes to the broader challenge of developing spatial models that are both mechanistically grounded and practically useful for understanding and predicting infectious disease dynamics, especially when dealing with multiple co-circulating strains in endemic settings. Furthermore, acknowledging the well-established mathematical correspondence between the classical Levins metapopulation model (Levins, 1969) and SIS phenomenology (Hanski and Gilpin, 1991; Hanski, 1991; Malagon et al., 2025), our results here on a multi-strain multi-patch SIS system with co-colonization, strain interactions and patch heterogeneity, can be directly useful to a wide range of spatial, evolutionary and community ecology research questions, beyond infectious diseases.

## 2. The model - metacommunity of hosts across *P* patches and *N* co-circulating strains

### 2.1. The SIS model with coinfection and host migration across patches

We describe the model over a discrete set of *P* separate patches (Figure 1). The populations within each patch are assumed to be well-mixed (i.e., no within-patch spatial or contact heterogeneity). Susceptible individuals (at density *S* ) are infected upon contact with infected individuals at a rate β. Single infected *I*^*i*^ can become co-infected upon further contact with infected individuals, but with an altered susceptibility coefficient, relative to a susceptible host. We allow for coinfection by the same strain (*I*^*ii*^) as well as by different strains (*I*^*ij*^), following the structural and evolutionary arguments in earlier coinfection models (van Baalen and Sabelis, 1995; Alizon, 2013). In particular, when a host is singly-colonized with strain *i*, it can become co-colonized with strain *j* with an altered susceptibility factor *k*^*ij*^. These altered susceptibility parameters between strains follow the earlier formulation (Madec and Gjini, 2020), and are introduced in the form of a matrix of coefficients *k*^*ij*^, which when above 1 indicate facilitation between strains *i* and *j*, and when below 1, indicate competition. Single infected individuals and coinfected individuals recover from infection back to the susceptible class at equal global rate γ, but there can be differences in clearing infection by each strain γ^*i*^, or composite infections γ^*ij*^ (for coinfection). There is no virulence so population size remains constant, with susceptibles having recruitment rate *r*, equal to the population per capita death rate. Note that all variables and parameters are to be understood as P-dimensional column vectors, with each entry being associated to one of the patches, labelled via *p* ∈ {1, ..*P* }. With this, the equations of the model, describing host proportions in different compartments, translated to discrete P-dimensional space, are:

**Figure 1:**
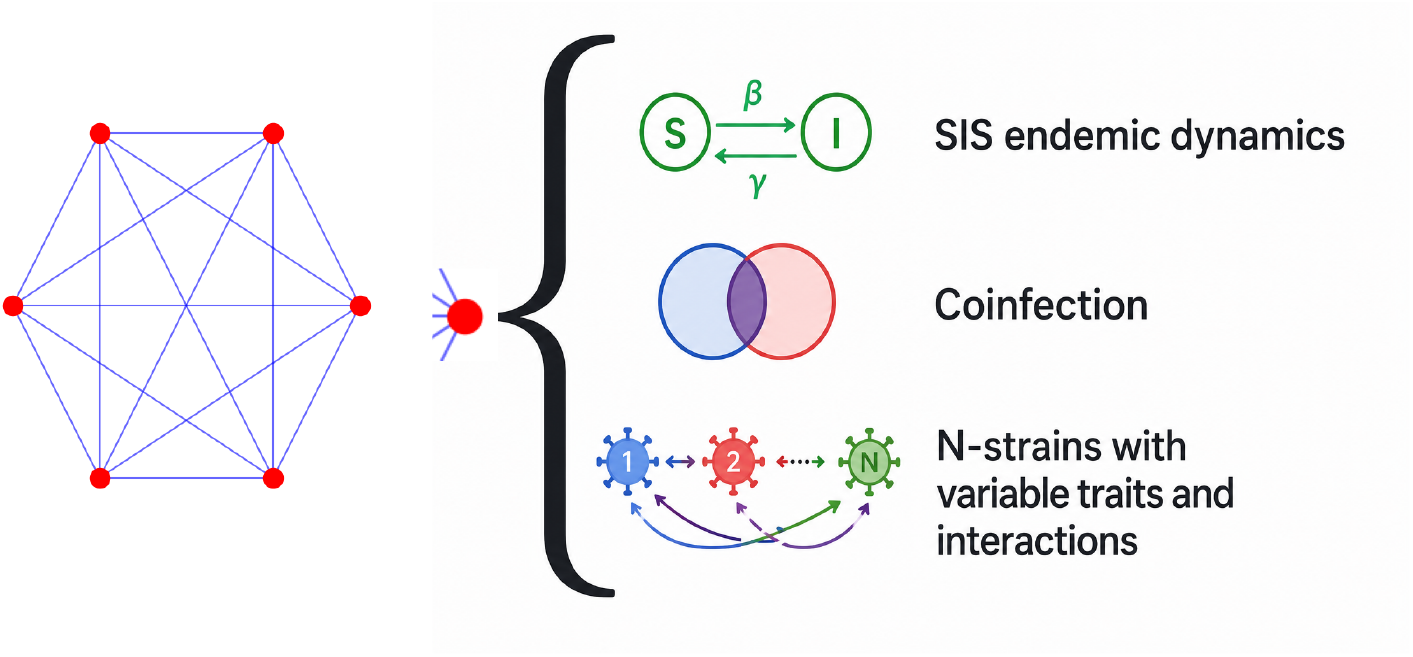
Schematic of multi-patch SID model structure with *N* strains. We model a metacommunity of hosts among a set of interconnected patches. In each patch SIS dynamics with coinfection happens among hosts, where *N* strains potentially co-circulate. The connectivity matrix between patches allows hosts to migrate at different rates in principle among any two sites, with each network edge having its own weight *d*_*pk*_, although our main simulations here assume the simplest case of uniform connectivity. All the patches may differ in quality, allowing for different mean-field parameters and strain fitness differences, as captured by the fast-slow dynamic approximation (Sec.3.1).

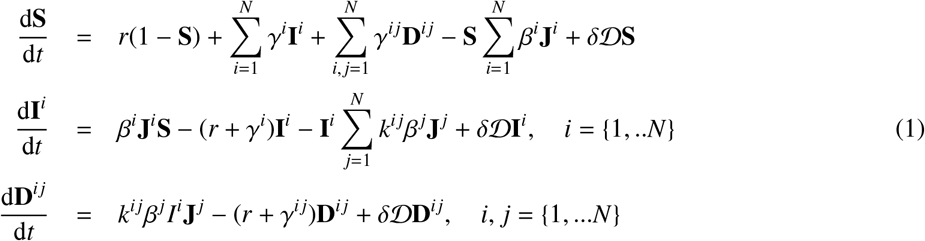

with D = (*d*_*pk*_)_*p,k*_ ∈ ℝ^*P*×*P*^ a Metzler matrix relating the connectivity between patches and δ the overall migration rate.

For the total population in each patch to be constant (and renormalized to 1) we assumed that D(1 · · ·, 1)^*T*^ = (0, · · ·, 0)^*T*^ which reads for all *p* ∈ ⟦ 1, *P* ⟧:

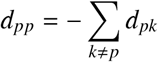

The force of infection for each strain *i* in a patch *p* is given by 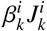 where the variable:

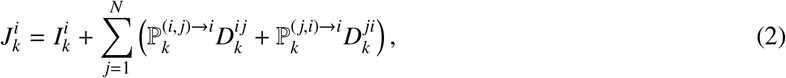

sums the contribution of all hosts (singly-and doubly-infected) that can transmit strain *i* in patch *k*, and where the strain transmission bias parameters for mixed coinfected hosts satisfy: 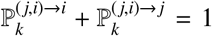. The total single infection prevalence in patch *p* is given by 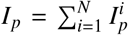. The total co-infection prevalence in patch *p* is given by 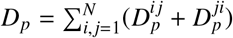. The total infection prevalence in patch *p* is given by *T*_*p*_ = *I*_*p*_ + *D*_*p*_.

In this model, which has of the order of (*N* + *N*^2^) × *P* equations, it is very hard to perform any analysis and anticipate the selection dynamics between strains. Furthermore, it is very difficult to quantify and disentangle the specific role of inter-patch heterogeneity, inter-strain diversity and host movement. To make the multi-strain multi-patch dynamics tractable and enable some analytic insights, we must adopt some simplifying assumptions. For this, we resort to the same simplifying assumptions of the analogous model considered in continuous space (Le and Madec, 2023). In this model, strain dissimilarity is modelled in the spirit of singular perturbation theory, through a small parameter ε, which applies to all strain-specific parameters of the model, expressed as a reference plus an ε −close deviation from that reference. This formulation together with the biological meaning of all parameters is presented in Table 1.

**Table 1:**
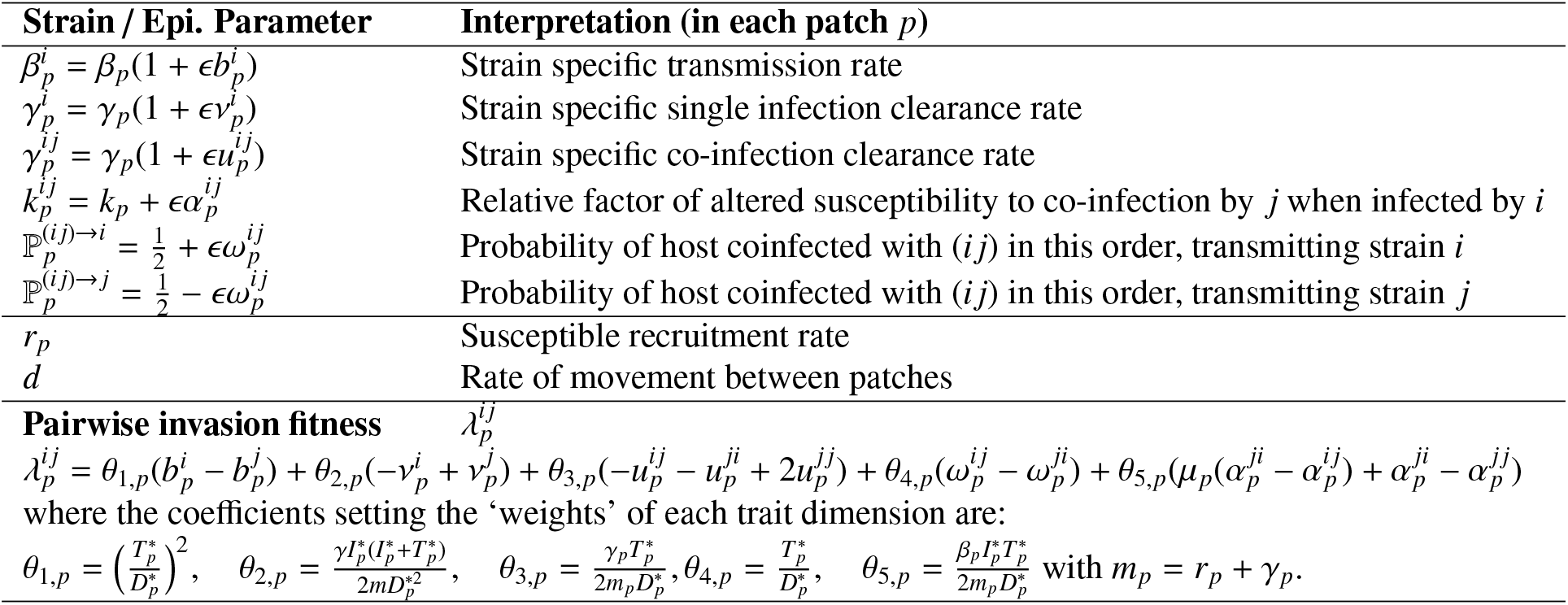
Strain similarity along 5 biological parameters. Notice that the model is general enough to encapsulate variation between patches both in the neutral model parameters β, γ, *k, r, m* (environmental mean-field heterogeneity) and in the non-neutral parameters 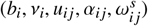, namely the relative variation between strains. However most of the time, we will assume explicitly only environmental heterogeneity between patches and study the net variation between strains across patches in terms of 4 effective λ coefficients, as shown in Eq. 13.

Like for the spatially continuous version of this model (see (Le and Madec, 2023)) we focus on a particular regime of the strain-dependent parameters and on the migration speed δ.

Let 0 < ε ≪1 be a small parameter. We require two conditions for the model reduction, based on similarity arguments, to work. These are:

#### H1 Quasi-Neutral assumption for co-circulating strains

In each patch *p*, the strain-dependent parameters are assumed to be ε-close. For instance, for the transmission rate of strain *i* in patch *p* we write 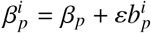, with 0 < ε ≪1 small. See Table 1 for the notation on all the parameters. Note that this is an assumption of similarity between strains. There is no assumption of the variability of the parameters between patches.

#### H2 Slow migration assumption

We assume that the migration rate of hosts is ε-small: δ = ε*d* for a fixed *d* > 0.

Note that it is essential that the same ε appears in H1 and H2. H2 essentially means that the host migration is slow compared to the global epidemic speed (typically driven by *R*_0_.) The parameter *d* is fixed but completely free. For instance large *d* means that movement is at an intermediate speed between the strain similarity (scale ε) and the global epidemiology.

## 3. Results

### 3.1. Discretization of the slow-fast approximation: explicit N strain-frequency dynamics across P patches

In the spatially continuous system, under the assumption H1 and H2, through a slow-fast decomposition, it has been shown in (Le and Madec, 2023) that, as ϵ →0 each aggregated epidemiological variable in each point in space approaches the equilibrium set by the neutral model (ϵ = 0, i.e. all strains equivalent):

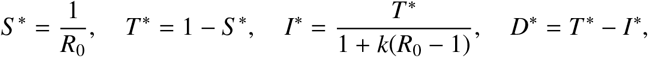

for susceptible, single-infected and co-infected host prevalence, with 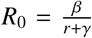 as the basic reproduction number. Note that these aggregated variables are indeed space-dependent. The model relies on *R*_0_(*x*) > 1 in each point in space, i.e. the persistence of colonization is endemic everywhere.

For the slow-variables *z*^*i*^, Le and Madec (2023) arrived at the following PDE equation with advection and diffusion, denoting the rates of change in strain frequencies over space and time *z*^*i*^(*x, t*):

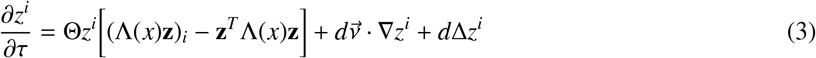

where the *z*^*i*^ represent the frequencies of strain *i* in each point in space, 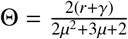 with 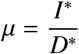 and

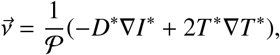

with 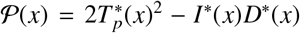 also space-dependent. Hence the vector 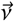 affecting the advection term in the local frequency equation (3), keeps a trace of the heterogeneity over space in mean-field transmission and other epidemiological parameters.

We will now discretize the spatiotemporal equations in the case of *N* strains with slow diffusion δ = ϵ*d* with ϵ ≪ 1, considering host movement among *P* patches. During discretization, the continuous space variable *x* is replaced with the integer *p* ∈ ⟦1, *P*⟧. This way, *z*^*i*^(*t, x*) is replaced with 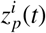, which gives the value of the frequency *z* for strain *i* in patch *p* (the *t* generally being implicit).

#### Patch-specific global epidemiological variables

Here we present the discrete version of the aggregated variables, ∀*p* ∈ ⟦1, *P*⟧, when *t* → +∞:

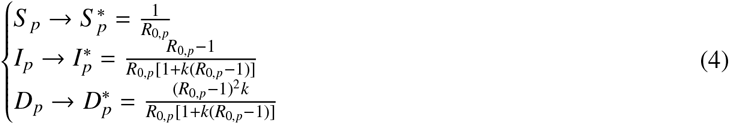

From these global variables, we also have several other key patch-specific quantities: ∀*p* (see Table 1), summarizing the ‘environmental quality’ of each patch.

#### Patch-specific selection forces

Now we can define the patch-specific pairwise invasion fitnesses between strains: ∀*i*, ∀ *j*, ∀*p*,

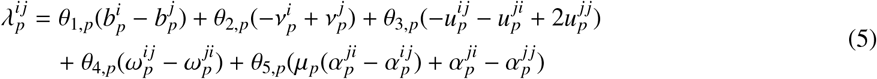

which give rise to the matrix 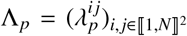 (local pairwise invasion fitness matrix among all strains *N* × *N*, experienced in patch *p*). Notice that these invasion fitnesses are weighted averages of strain variation along each of the 5 key dimensions (Table 1), like in (Le et al., 2023; Le and Madec, 2023).

Considering the connectivity matrix D between all patches, we can write the correspondence between the continuous replicator model and the discrete replicator equation: ∀*p* ⟦∈ 1, *P*⟧ and ∀ *i*∈ ⟦1, *N* ⟧following the steps in Table 2.

**Table 2:** Elements of the discretization of the PDE system, from continuous space to discrete space.

| PDE element | Discretized element |
| --- | --- |
| $z^i(x)$ | $z_p^i$ |
| $\mathbf{z}(x) = (z^i(x))_i$ | $\mathbf{z}_p = (z_p^i)_{i \in [1, \dots, N]}$ |
| $z^i : x \mapsto z^i(x)$ | $\mathbf{z}^i = (z_p^i)_{p \in [1, \dots, P]}$ |
| $(\Delta z^i)(x) = \text{div}(\nabla z^i)(x)$ | $\sum_{k=1}^P d_{pk}(z_k^i - z_p^i) = (\mathcal{D}\mathbf{z}^i)_p$ |
| $(\nabla z^i)(x)$ | $\left[ \sqrt{d_{pk}}(z_k^i - z_p^i) \right]_k$ |
| $\vec{v}(x) = \frac{2T^*(x)\nabla T^*(x) - D^*(x)\nabla I^*(x)}{\mathcal{P}(x)}$ | $\frac{1}{\mathcal{P}_p}(v_{pk})_k = \frac{1}{\mathcal{P}_p} \left( \sqrt{d_{pk}}(2T_p^*(T_k^* - T_p^*) - D_p^*(I_k^* - I_p^*)) \right)_k$ |
| $(\Delta + \vec{v}(x) \cdot \nabla)(z^i)(x)$ | $\sum_{k=1}^P d_{kp}(1 + v_{kp})z_k^i = (\mathcal{M}\mathbf{z}^i)_p$ |

Therefore, the discretization of (3) shows that the dynamics of the frequency 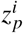 of strain *i* in patch *p*, within a multi-patch framework, is given by:

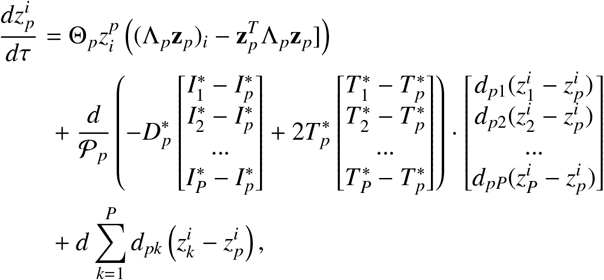

which can be simplified as:

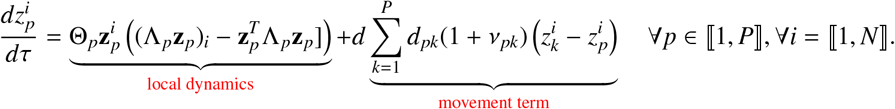

In the above spatial replicator equation, we have that the patch heterogeneity explicitly modulates the net contribution of host movement to strain dynamics, via the parameters ν_*pk*_: ∀*p*, ∀*k*

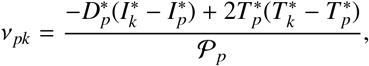

measuring the relevant epidemiological (mean-field environmental) ‘gradient’ between any two patches *p* and *k* in the system. The speed of local dynamics Θ_*p*_ is also patch-specific (see Appendix A for more details on these parameters). Note that we have

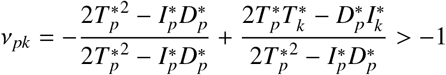

the later inequality coming from the fact that 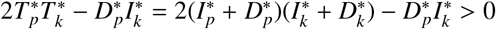 for any *k*.

Let us denote 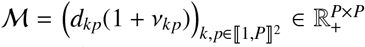 the net connectivity matrix whose coefficients measure the effective strength of links between patches, after accounting for inter-patch mean-field heterogeneity. The fact that 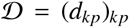 is an irreducible Metzler matrix and that 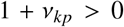 imply that *M* is also an irreducible Metzler matrix. With these notations, the multi-strain dynamics system reads compactly, for all strains *i* = 1, ..*N* and patches *p* = 1, ..*P*:

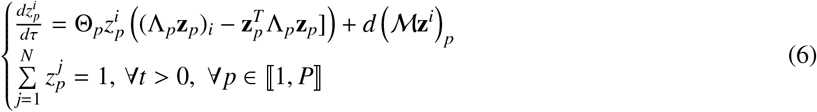

This is the system that describes local strain dynamics explicitly in each patch, and that we study in the following sections.

#### Population-level dynamics in each patch

An immediate consequence of this fast-slow approximation framework for the SIS model with coinfection, similar to (Madec and Gjini, 2020; Le and Madec, 2023), is that the final epidemiological compartmental variables in each patch *p* can be obtained after strain frequencies as:

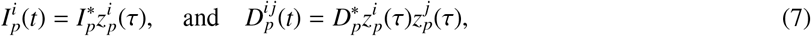

for single and co-infection respectively, which in our case will be patch-specific, first because the global prevalence compartments (*S* _*p*_, *I*_*p*_, *D*_*p*_) can be different between the patches (Eqs. 4), secondly because the strain frequencies 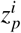 can be different between the patches (Eq. 3).

#### Much more than a model reduction

Thus we have arrived at a substantial model reduction of dimension (*N −*1)*P*, starting from the original population structure system which has of the order (*N* + *N*^2^)*P* equations. Here, we have presented the path of discretization of the PDE (Le and Madec, 2023), because it is easier to follow, but the same final model reduction (6)-(7) can be formally obtained by deriving explicitly the slow-fast approximation and its regularization, directly from the SIS multi-strain multi-patch ODE system (1). For a complete bottom-up derivation of such slow-fast approximation, see companion paper (Madec and Gjini, 2025). Besides the consistency of the two derivation approaches, this new formulation of the multi-strain dynamics as a replicator system modulated by movement and inter-patch heterogeneity makes it both computationally and analytically easier to study the role of different parameters and factors on the ultimate quality of coexistence of interacting strains. Furthermore, encapsulating inter-strain relative fitness and inter-patch heterogeneity into a few relevant aggregated parameters substantially enhances the robustness of eventual parameter inference from such dynamics in empirical data. Next, we will illustrate some special cases of this model to highlight insights that can be obtained analytically on endemic SIS systems from this backbone formulation, and emphasize that even in low-dimensional cases, understanding strain selection processes among heterogeneous patches is hard.

### 3.2. Some general model properties

#### Defining global strain fitness for general N and P

When *d* = 0, meaning that no migration occurs between patches, each patch behaves as an independent epidemiological system. In this case, the global evolutionary dynamics reduce to a set of uncoupled local competitions, each governed by its own invasion matrix Λ_*p*_. The pairwise invasion fitnesses 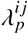 defined in (5) then have a direct interpretation: 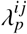 is the invasion rate (in the time unit 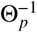) of strain *i* within patch *p* when that patch is occupied exclusively by strain *j*.

For positive *d*, the patches are coupled and this local interpretation no longer holds. Instead, there exists a *global* invasion rate ρ^*ij*^ = α_max_(A^*i j*^), defined as the maximal eigenvalue of the irreducible Metzler matrix

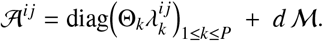

The quantity ρ^*ij*^ admits the following interpretation: if strain *j* is the only strain present in the system (in every patch), then ρ^*ij*^ is the global growth rate of strain *i*. In particular, strain *i* can invade strain *j* if and only if ρ^*ij*^ > 0.

Hence, this global invasion rate is the key classical quantity enabling the use of standard mathematical tools as well as modern ecological theory, such as Chesson’s coexistence framework Chesson (1994) or invasion graph theory Hofbauer and Schreiber (2022). It is worth noting that such a quantity is not straightforward to define directly from the full multi-strain system (1). Our reduction therefore has the important consequence of making classical spatial ecology methods applicable to multi-strain epidemiological systems.

#### The case of some identical patches

By identical patches, we mean the same in all aspects. Two patches are called identical if they verify the same initial conditions, have the same parameters and strain species. It is an interesting case because it allows to perform a system reduction. If patches 1, 2, …*m* are identical, (for *m* ∈ ⟦2, *P* ⟧) from 6 it remains only to solve: ∀*p* ∈ {*m*, · · ·, *P*}

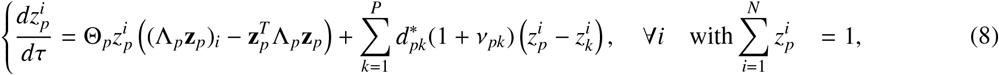

and with

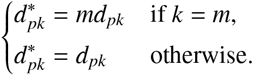

And the *m* − 1 first patches follow the same dynamics as patch *m* : *i, z*_*i*,1_ = *z*_*i,m*−1_ = *z*_*i,m*_. This shows that the *N*-strain *P*-patch network with *m* equal patches is equivalent to an *N*-strain (*P − m*)-patch reweighted network, allowing to reduce problem dimension.

In particular, when all patches are equivalent, the system reduces to the single-patch situation, and for every patch *p* the global *i* → *j* invasion rate is given by:

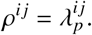

This shows that the dynamical phenomena induced by space and impacting strain coexistence —such as invasion outcomes—are essentially driven by *heterogeneity* between patches, and not by migration alone.

#### Comparison with the non-spatial replicator

It is clear that the role of ν_*pk*_, and the role of *d*_*pk*_ are crucial for the difference of Eq. (6) with the non-spatial model, or single-patch model. The equations for focal strain frequency in each patch have now two clear components: a first component, which is just the classical replicator dynamics (purely local selection dynamics in each patch), and a second component, which is the net benefit from global movement experienced by the focal strain in the current patch and in the rest of the patches (net metapopulation effect).

While the speed parameter Θ is inconsequential in the single-patch model for determining final selection outcome, in the coupled P-patch system, the speed of the dynamics in each patch matters for final selection outcome: Θ_*p*_ determine often both the magnitude and stability of the equilibria. Another immediate thing to notice is that under equal neutral *mean-field* parameters between patches, there is no global spatial heterogeneity in the system, ν_*pk*_ = 0 for all *p, k*, and Θ_*p*_ = Θ, so the contribution of host movement to strain dynamics in that case is purely driven by diffusion.

Regarding the complexity of dynamics, we can say that while in the single-patch system, the magnitude of the unique coexistence equilibrium is very easy to compute (see for example Table 3 for *N* = 2) and its stability can be unambiguously decided on the basis of studies of the marginal exclusion equilibria (**0** = (0, · · ·, 0) and **1** = (1, · · ·, 1)), in the coupled *P*-patch system, the coexistence equilibria have no straightforward analytic solutions, there can be multiple such interior coexistence equilibria, and hence the stability of interior coexistence points cannot be straightforwardly decided from studying simply the behaviour of the marginal exclusion equilibria **0** and **1**. Even though analysis of this spatial system is harder, we can still advance theoretically and computationally in some special cases, and derive explicit criteria. Certainly, the analytical tractability is much more within reach via this replicator formulation (6), than by addressing the full epidemiological system (1).

**Table 3:**
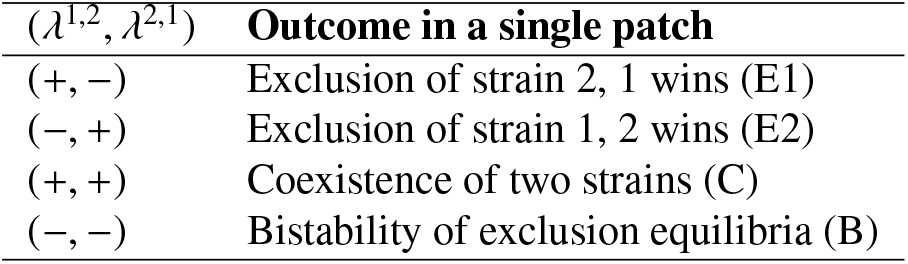
The final ecological outcomes for *N* = 2, in each patch, when uncoupled, depend purely on the mutual invasion fitnesses between 2 strains.

| $(\lambda^{1,2}, \lambda^{2,1})$ | Outcome in a single patch |
| --- | --- |
| (+, -) | Exclusion of strain 2, 1 wins (E1) |
| (-, +) | Exclusion of strain 1, 2 wins (E2) |
| (+, +) | Coexistence of two strains (C) |
| (-, -) | Bistability of exclusion equilibria (B) |

##### Box 1. Results on the general *N* = 2 strains, *P*-patch model

When *N* = 2 we have 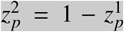 . Hence, denoting 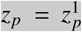 and **z** = (*z*_*p*_)_*p*_, the system (6) reads shortly for *p* ∈ ⟦ 1, *P* ⟧

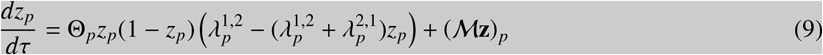

We have 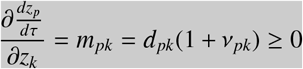 for any *p* ≠ *k*. Then the system is cooperative. By the theorem 1.1 p.56 in Smith (1995), the irreducibility of *M* implies that the system is strongly monotonous (see theorem 1.1 p.56). As a consequence, in the particular case of *N* = 2, we obtain some general properties that are inherited from the strongly monotone structure. This fact leads to several important properties. We refer to Smith and Waltman (1995), where Chapter 6 presents results closely related to our current setting, and Appendix C in this book, provides a concise rationale for the properties of monotone systems. For further details on monotone systems, see Smith (1995). Without being exhaustive, let us now state 3 key properties of system (9).

(i) There are no attracting limit cycles.
(ii) Nearly all the trajectories converge to a steady state. To state the last properties, we note the simple fact that the trivial solutions **0** = (0, …, 0)^*T*^ ∈ ℝ^*P*^ and **1** = (1, …, 1)^*T*^ ∈ R^*P*^ are both stationary solutions of the system (9), corresponding respectively to 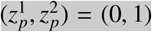 for all *p*, and 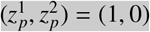for all *p*.
(iii) If both **0** and **1** are linearly unstable, then there exist two steady states **z**^∗^ and **z**^∗∗^ satisfying 0 < **z**^∗^ **z**^∗∗^ < 1 such that the set 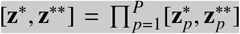 attracts any trajectory **z**(*t*) with initial state **z**(0) ∈ [0, 1]^*P*^\{ **0, 1** }. In particular, if **z**^∗^ = **z**^∗∗^ then this coexistence solution is unique and globally stable.
(iv) If both **0** is unstable and **1** is stable then either their exists a non trivial coexistence solution or **1** is globally stable.

Similar results hold under different stability assumptions on **0** and **1**. See Smith and Waltman (1995), Chapter 6, for examples of such results in a similar framework.

It is important to highlight the fact that the strong monotonicity of this system essentially allows one to reduce the study of its dynamics to the mere existence and stability of steady states.

However, even though this reduces the complexity of the analysis, due to spatial heterogeneity, an exhaustive study of system (9) remains out of reach. This is why we restrict our study to the cases of *P* = 2 and *P* = 3 patches.

#### 3.3. Some theoretical results for the multi-patch system for N = 2

##### A sufficient condition of competitive exclusion for N = 2 strains and any number of patches P

As explained in Box 1, for *N* = 2 the system is strongly monotone, which is a powerful property. In this part, we focus on the case where the same strain wins in competitive exclusion in each patch.

Let’s assume that without migration, strain 1 is dominant in each patch, that is:

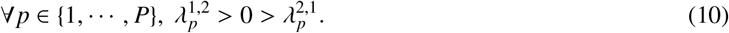

We will show that in that case **1** is globally stable.

Indeed, under assumption (10), it is easy to show that **0** is unstable and that **1** is stable. By point (*iv*) of Box 1, in order to prove that **1** is globally stable, it suffices to show that there is no non-trivial (interior) equilibrium.

The system reads

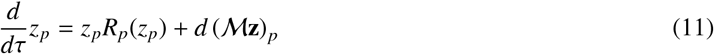

where we denote

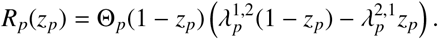

We proceed by contradiction. Assume that there exists a coexistence solution **z** ∈ (0, 1)^*P*^. Then **z** is a positive solution of

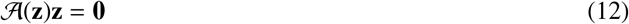

wherein we have set *A* (**z**) = *d M* + diag(*R*_*p*_(*z*_*p*_)). Like the matrix *M*, the matrix *A*(**z**) is an irreducible Metzler matrix.

Firstly, (12) shows that **z** ∈ (0, 1)^*P*^ is a positive eigenvector of *A*(**z**) for the eigenvalue 0. By the Perron-Frobenius theorem for Metzler matrices (see Bullo (2024), Theorem 10.2), this implies that 0 is the principal eigenvalue of A(**z**). Secondly, we also have *M***1** = **0**, which yields

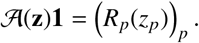

From assumption (10), we see that *R*_*p*_(*z*_*p*_) < 0 for each *p*, and thus *A* (**z**)**1** < 0. By Theorem 14.1 (iv) of Bullo (2024), this implies that the principal eigenvalue of *A* (**z**) is negative, which contradicts (12). This proves that when the same strain wins in competitive exclusion in each patch, it is the global winner also in the multi-patch system.

## 4. Coexistence and stability: easy to analyze for special cases of *N* = 2 strains and 2 or 3 patches

The case of 2 co-circulating strains is the easier one to tackle analytically (Box 1) and often quite relevant for applications (e.g. focal and non-focal pathogen, antibiotic-resistant vs. sensitive strain, wild type vs. mutant variant), so here, we will focus on this case in a metapopulation composed of 2 or 3 patches, or environments, where hosts can move. We will generally assume in this work that movement is uniform across patches, *d*_*pk*_ = *d* for all *p, k*, however the model is general and encapsulates cases beyond this symmetric movement assumption. Regarding patch heterogeneity arising i) from mean-field global parameters (ν_*pk*_ ≠ 0, Θ_*p*_ patch-specific), and from ii) patch-specific strain invasion fitnesses (Figure 2), a natural question that arises and that we will address is the following:

**Figure 2:**
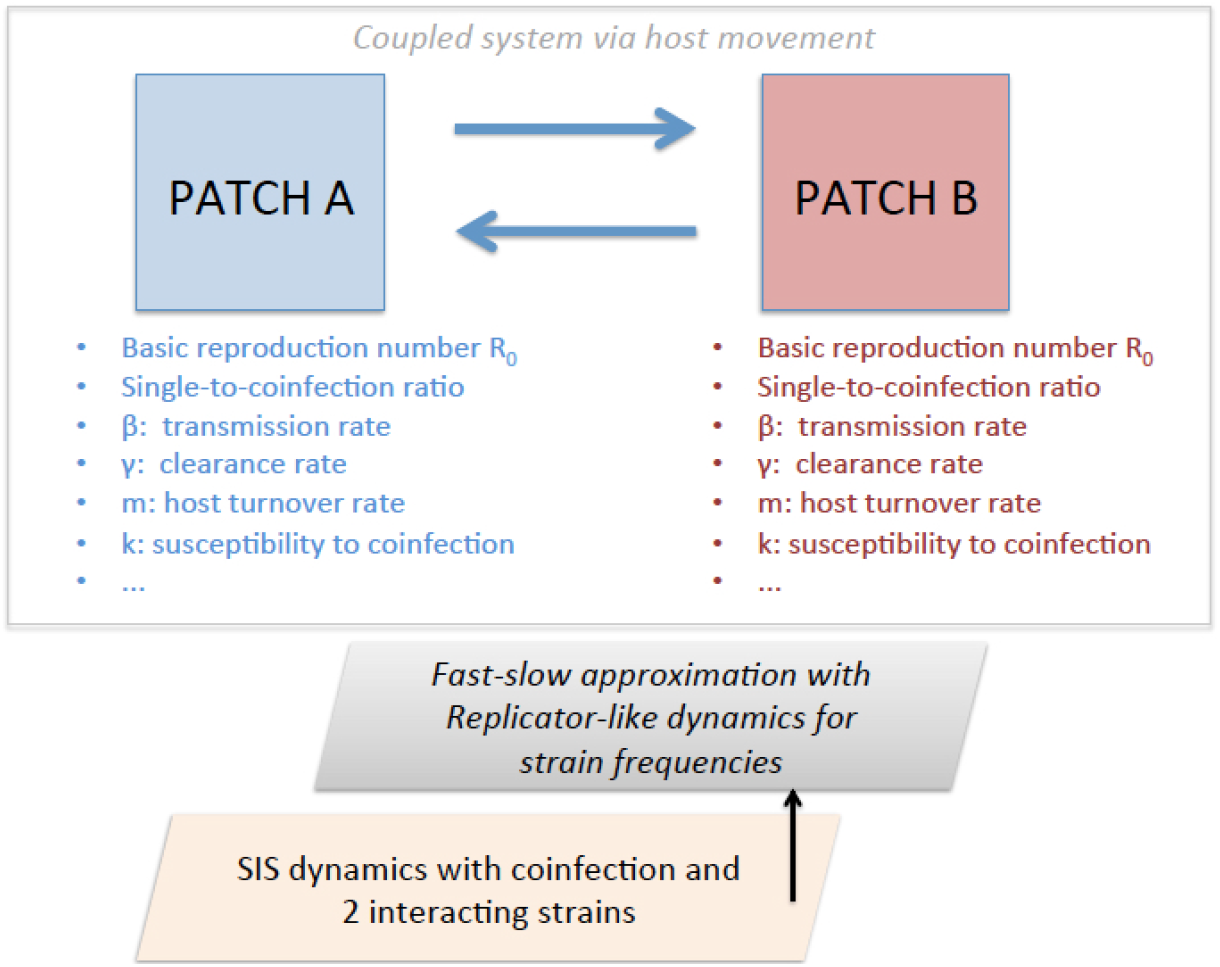
Schematic illustrating the context of the epidemiological SIS coinfection model for *N* = 2 strains and *P* = 2 patches. We show an epidemiological model with two patches *A* and *B* and 2 interacting strains, where hosts move in space and follow broadly SIS dynamics with coinfection. Assuming strain similarity,following (Le and Madec, 2023) we can decompose the dynamics into a fast and slow component, and consider the special case of the reduced model in the limit relatively slow diffusion. In the discrete patch case, we arrive at a replicator-like equation for strain frequency in each patch. Besides local selection, this equation contains explicitly the contribution of patch heterogeneity and global migration, to the competitive dynamics between strains.

*How do these different heterogeneity layers interplay with each other to shape strain selection in the metacommunity, and what is the role of host movement?*

### The model for P = 2 patches and N = 2 strains

Writing the spatial replicator equation (6) in terms of the frequencies of one of the strains (e.g. strain 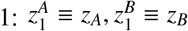), and for the case of 2 patches, we obtain:

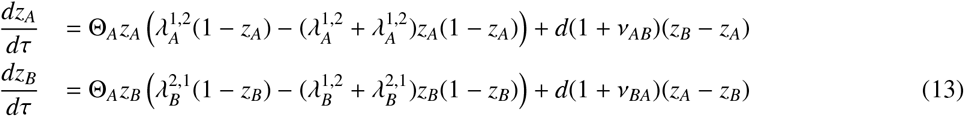

where we add the subscripts A and B, to the strain invasion fitnesses in patch A and B, respectively.

Although the equations governing strain selection dynamics across two patches look very simple, they can generate very complex behavior, - where up to nine possible equilibria are possible - unlike the single-patch system. Our scope is not to provide an exhaustive exploration of all cases, but rather to illustrate a few key scenarios where the coupling between patches can unequivocally favour exclusion or coexistence, and maintain or overturn the final outcome predicted by each single patch alone (Table 4). We will focus only on analysis of selection dynamics between strains, given fixed parameters (Eq. 3), as subsequent translation to epidemiology is straightforward via Eq.7.

**Table 4:** Possible outcomes for the ecological dynamics of 2 strains circulating across two heterogeneous patches connected via host movement. Outcome relative to a focal strain (here strain 1) frequency in patch A and patch B. We denote by (0) and (1) the outcome of single-patch dynamics favouring exclusion of strain 1 or persistence of strain 1. We denote by (0,0), (1,1) the case where the coupled patch dynamics favours competitive exclusion or competitive dominance of strain 1 in both patches. The scenarios of strain coexistence in single patch or in both patches are denoted by (*z*) and (*z*_*A*_, *z*_*B*_) respectively.

| | **Patch A**<br>$(\lambda_A^{1,2}, \lambda_A^{2,1})$ | **Patch B**<br>$(\lambda_B^{1,2}, \lambda_B^{2,1})$ | **Without movement, A and B**<br>Uncoupled outcome in each patch | **With movement $A \leftrightarrow B$**<br>System outcome after coupling |
| --- | --- | --- | --- | --- |
| **1.** | (+, -)<br>(-, +) | (+, -)<br>(-, +) | (1), (1)<br>(0), (0) | (1, 1)<br>(0, 0) |
| **2.** | (-, +) | (+, -) | (0), (1) | (1, 1)<br>(0, 0)<br>$(z_A, z_B)$<br>multistability: (1,1),(0,0) or more |
| **3.** | (+, +) | (+, -) | (z), (1) | $(z_A, z_B)$<br>(1, 1)<br>multistability, excl. (0,0) |
| **4.** | (++) | (+, +) | (z), $(z')$ | Permanence, $(z_A, z_B)$, multistability possible, |
| **5.** | (-, -) | (-, -) | (0/1), (0/1) *bistable-bistable* | (1,1), (0,0), $(z_A, z_B)$<br>always multistability: (1,1) and (0,0) or more |

−

#### Coupling (1) (1) patches

Without loss of generality, this is the simplest case to analyze, (equivalent to (0)-(0) coupling). In this case the same strain is a competitive single winner in either patch when uncoupled (monostability of same exclusion in each patch). By the general result in 3.3; (1, 1) is globally stable.

#### Coupling (z) − (z^′^) coexistence patches

This is also a simple case to analyze. In this case the two strains coexist in either patch when uncoupled (stability of of coexistence in each patch). This means both strains have positive individual invasion fitnesses in each patch: 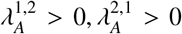 in patch A,and 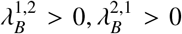 in patch B. It is straightforward to verify that that in this case, the two marginal equilibria (1,1) and (0,0) are both unstable, hence the only stable equilibrium of the system, independently of other patch-specific parameters, must inevitably be some interior coexistence point 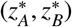. Imposing the positive determinant and negative trace stability criteria on the Jacobian evaluated at each marginal equilibrium, leads to a contradiction, hence proving the impossibility of such scenario under this parameter regime.

#### Coupling (0) − (1) patches: patch heterogeneity, invasion fitnesses and migration rate drive final outcome

This is the most interesting case. Each patch alone favours the exclusion of a different strain. For illustration, we will study here a simplified scenario, assuming the invasion fitnesses are fixed free parameters and do not depend on mean-field parameters of the system. We assume an arbitrary fixed Λ matrix guaranteeing these patch-specific regimes. We examine numerically the stability of the two marginal equilibria (0,0) and (1,1) in the coupled system.

Fixing values for λ^*i, j*^ in each patch, and migration parameter *d*, we can broadly assume the following relations for epidemiological parameters between patches: *R*_0,*B*_ = *cR*_0,*A*_ and µ_*B*_ = ωµ_*A*_, allowing to vary relative heterogeneity (see Appendix A for detailed parameter consequences). Under a certain value of *c* we can plot resulting qualitative regions of stability in the 2-patch system on the (*R*_0_, µ)-plane, based on evaluating the stability properties of the marginal equilibria (0, 0) and (1, 1) (Figure 3). Each region represents one of the possible outcomes for the system at each point (*R*_0_, µ) which are the parameters of patch A. Thus, in figure 3 we observe how the coupled 2-patch system can arrive at four possible broad selection outcomes depending on the basic reproduction number of patch A, *R*_0_, and the ratio of single to co-colonization µ = *I*/*D* (here assumed equal between patches).

**Figure 3:**
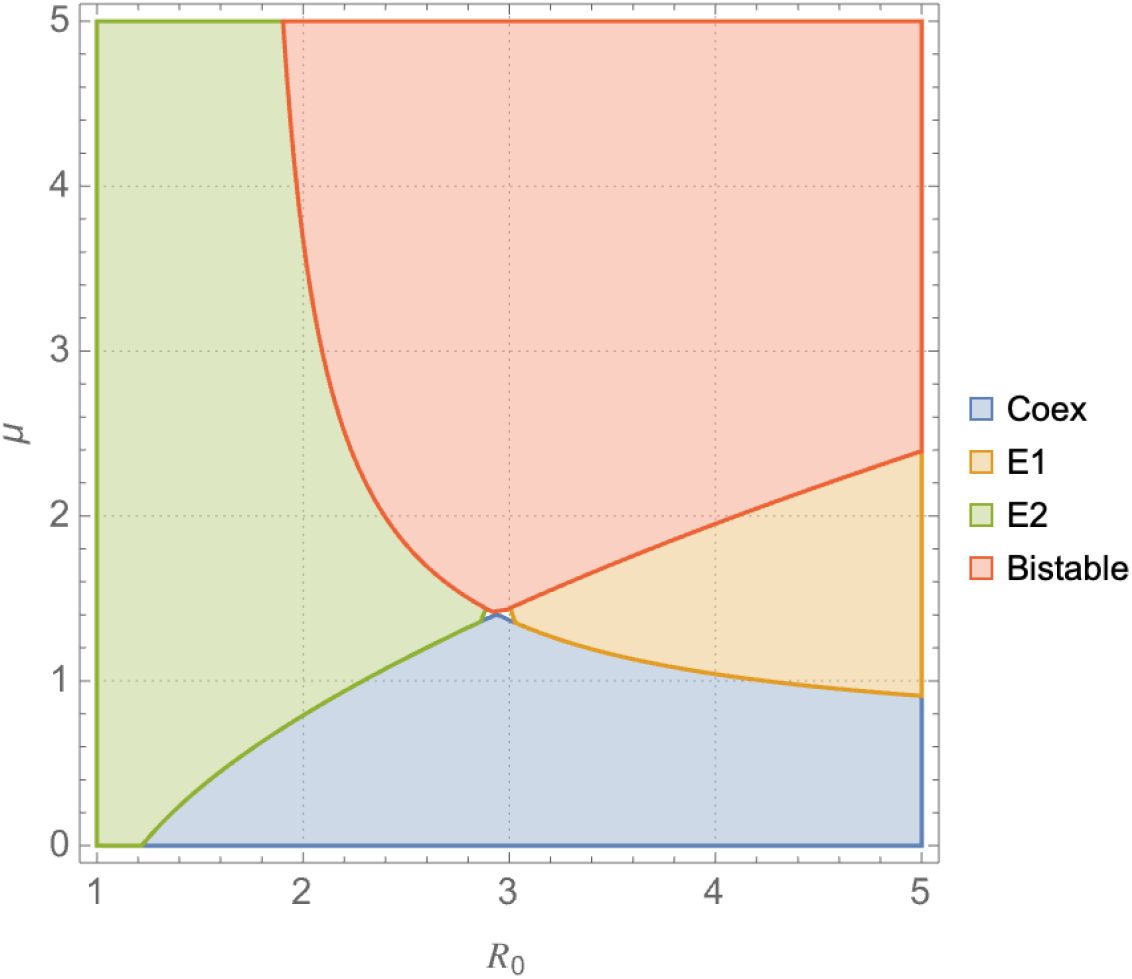
Outcome regions for the coupled 2-patch 2-strain system where alone patch A favours the exclusion of strain 2 and patch B the exclusion of strain 1, but patch B has higher total infection prevalence 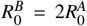. Patch A parameters: 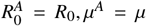 and strain invasion fitnesses 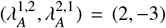. Patch B parameters: µ^*B*^ = µ and 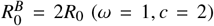, and invasion fitness coefficients 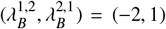. Migration rate between patches: *d* = 1, and demographic parameter for host turnover equal in each patch: *m* = 1.5. Notice that every time we change parameters of patch A, this changes also the parameters of patch *B*, despite the relative ratio between the two is held fixed. Inevitably, the precise shape of these regions (based on stability of marginal equilibria) does heavily depend on the choice of fixed parameters but they still provide good insight about the outcome of the system and can be used as a starting point for further numerical or analytical exploration. As a a function or patch variation in *R*_0_ and µ we can obtain different regimes of system behavior. And within each of these regimes, there is a potential for underlying complexity, such as multiple coexistence points and multi-stability of several equilibria. See also Appendix D.

For high values of the single-to-coinfection ratio, µ, coexistence of the two strains is impossible, and the 2-patch system always approaches an exclusion, be it mono-stable or bistable. The magnitude of *R*_0_, which scales the total infection prevalence, on the other hand, decides the winning strain. In figure 3, high values of *R*_0_ benefit strain 1, whereas low values benefit strain 2. What is interesting to notice here is that unlike the simple case of exactly anti-symmetric invasion fitnesses within each patch (full analysis in Appendix B), for arbitrary four λ coefficients across patches, bistability of (0,0) and (1,1) also becomes possible in the coupled system. In Appendix D, we also provide more details and numerical explorations of the case in Figure 3, including the effect of varying host movement rate *d*. In this particular example, increasing host movement rate, reduces the region of 2-strain coexistence in µ − *R*_0_ parameter space.

#### Connecting two originally bistable patches: the (0/1) − (0/1) case

This type of system naturally leads to more complex behavior, as the strains’ final outcome is frequency-dependent on initial conditions in each patch. In this case, it can be proven that the (0, 0) and (1, 1) global exclusion equilibria are both stable in the 2-patch system, implying the existence of unstable interior coexistence points, but without excluding the possibility of eventual stable coexistence points too. We illustrate such cases by plotting the nullclines of *z*_*A*_ and *z*_*B*_ for different assumptions of parameter variation between patches (Figures 4-5). These figures show that the resulting behavior of the system depends heavily on the global patch heterogeneity, be it in basic reproduction number *R*_0_ or single to coinfection ratio µ. In the *R*_0_ variation case (Figure 4), we see the most complex dynamics happens when *R*_0_ in patch B is 1.15 times higher than the *R*_0_ in patch A (middle scenario); there are 9 equilibria for this system, including 7 interior coexistence points. The marginal equilibria are still stable, however this does not guarantee global exclusion of either strain, because there are two interior coexistence points that are stable and could be reached depending on initial conditions. The other two values for *c* confirm the exclusive bistability of the two marginal equilibria.

**Figure 4:**
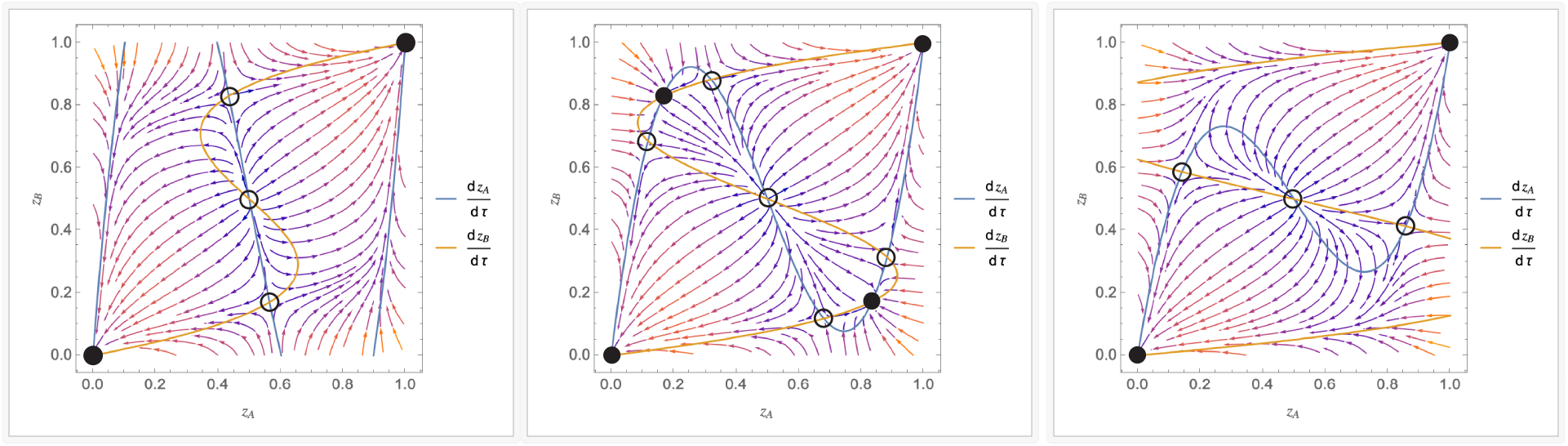
Connecting two bistable patches, under different *R*_0_ gradient. The parameter *c*, different in each plot, varies the *R*_0_ (transmission) gradient between patches, 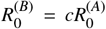. Left: *c* = 0.8 (2 stable equilibria) Middle: *c* = 1.15 ( 4 stable equilibria) Right: *c* = 2 (2 stable equilibria). In these plots the pairwise invasion fitnesses between the two strains are equal 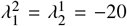 in each patch (equal ‘strength’ of bistability). The fixed parameter values are: 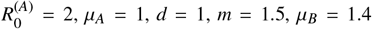. Stable/unstable steady states are denoted by the filled/empty circles, respectively.

In the coinfection gradient case (Figure 5), we also see a sensitivity to the relative magnitude of coinfection in each patch, where depending on parameter values, and initial conditions, bistability in each patch may drive ultimately interior stable coexistence everywhere in the system, instead of the classically expected exclusion.

**Figure 5:**
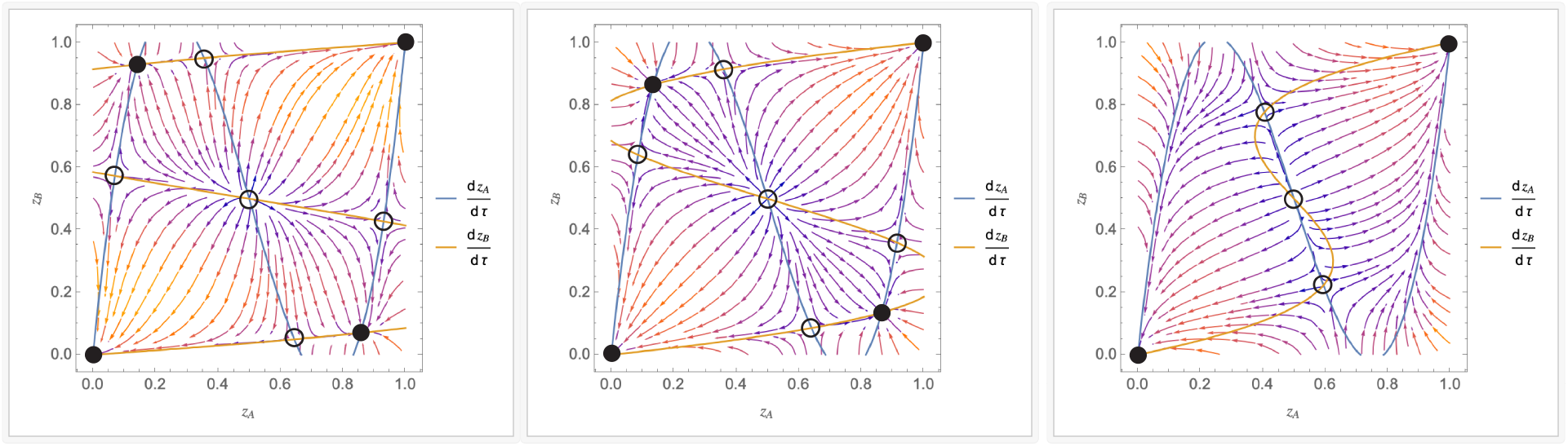
Connecting two bistable patches, under different coinfection gradient. The parameter *w*, different in each plot, varies the µ gradient between patches (ratio of single to co-infection) µ_*B*_ = *w*µ_*A*_, keeping *R*_0_ constant, here 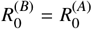. Left: *w* = 0.5 (4 stable equilibria) Middle: *w* = 1 (4 stable equilibria) Right: *w* = 2 (2 stable equilibria). In these plots the pairwise invasion fitnesses between the two strains are equal 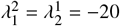 in each patch (equal ‘quality’ of bistability). The fixed parameter values are: 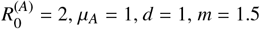.

In the scenarios above, we considered identical λ^*i, j*^ values in both patches, meaning the bistability in each original patch is of equal magnitude - this led to a certain symmetry in the phase-plots, visible in the intersection structure between the two nullclines. When we relax this symmetric stability assumption, and instead allow for the magnitudes of the λ’s to vary between strains and between patches, while still remaining negative, we see a broken symmetry in the corresponding phase-plots, and having 2 additional stable interior coexistence points becomes more improbable. In such a scenario, we have also checked the effect of host movement rate on the final outcome of the system (Figure 6). We see that in this asymmetric λ parameter regime and equal global epidemiological parameters, increased movement rate between patches reduces the chance strain coexistence, by reducing the number of alternative stable steady states in the system and pushing the system to the classical marginal equilibria: either one strain or the other everywhere.

**Figure 6:**
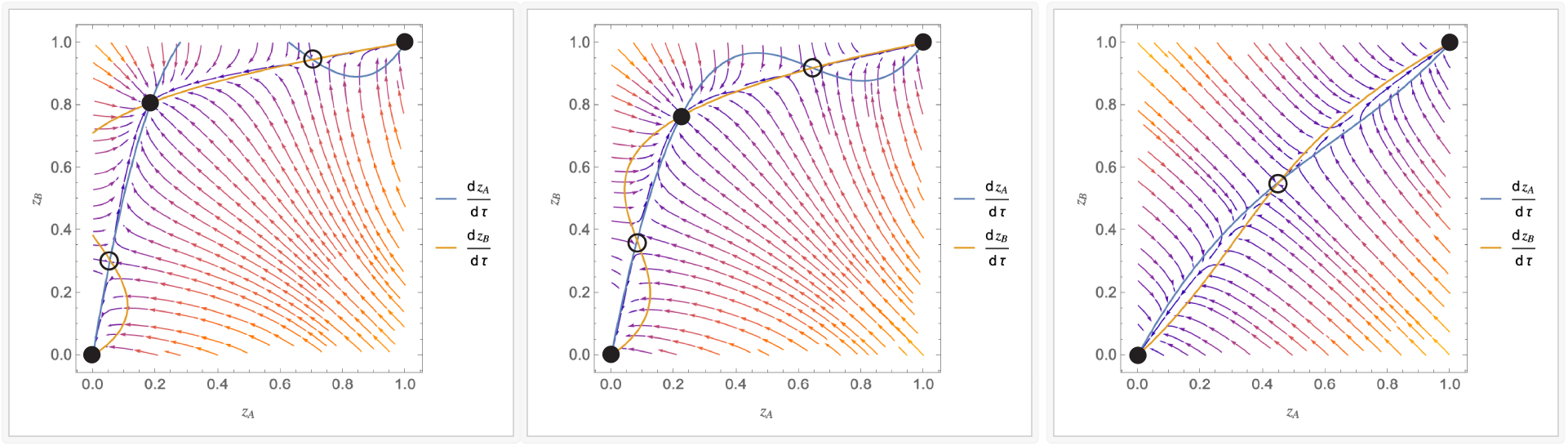
Effect of host movement rate *d* on strain dynamics between 2 locally bistable patches. Left: *d* = 0.8 (3 stable equilibria) Middle: *d* = 1 (3 stable equilibria) Right: *d* = 5 (2 stable equilibria). In these plots the pairwise invasion fitnesses between the two strains are unequal 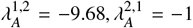 in patch A, and 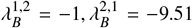 in patch B. The fixed parameter values are: 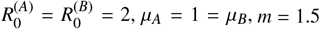 *m* = 1.5.

### 4.2. The model for N = 2 strains, P = 3 patches

This case is more complex to study, as there is one extra environment where the strains can compete for host resources, and this will affect their net fitness and dynamics in the system. In the 3-patch case, given by the following equations:

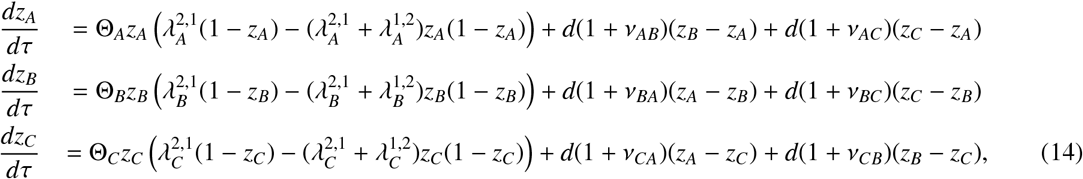

there appears to be the two usual trivial equilibria: (1,1,1) and (0,0,0), and potentially other coexistence ones, which are very complex to solve for since it comes to intersecting three cubics in 3D. Substitutions for 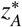 and 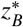 lead to finding the zeros of a degree 27 polynomial in 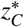, so there are at most 25 coexistence equilibria.

Despite this complexity and high-dimensionality, one can notice straightforward outcomes of the dynamics in special cases. For example, in the case where pairwise strain fitnesses are proportionally-related by the same positive constant in each patch, 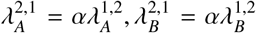 and 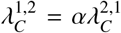 for some α > 0, the coexistence point 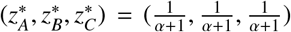 is also an equilibrium of the system. Another result is that mixed trivial outcomes are not possible, for instance (1,0,1) or (0,0,*z*_3_) because of the movement terms. Besides, some outcomes should be easily predicted. For instance, when connecting patches that have a stable exclusion steady-state on their own, coupled dynamics will lead to a global exclusion everywhere of that strain: (1), (1), (1) → (1, 1, 1) and (0), (0), (0) → (0, 0, 0), similar to the 2-patch case.

#### Three equal patches case

For the sake of simplicity and illustration, we consider here the case of equal patches, in the sense of the neutral model (Eqs. (4)). Some analysis can be performed in a straightforward manner (see Appendix C) and we can obtain a criterion for the stability of the marginal equilibria. Even if these results are partial, the stability of the exclusion state **0** and **1** says a lot about the dynamics thanks to the strong monotonicity of the system. Introducing aggregated parameters: 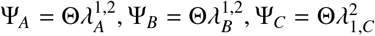, we have:

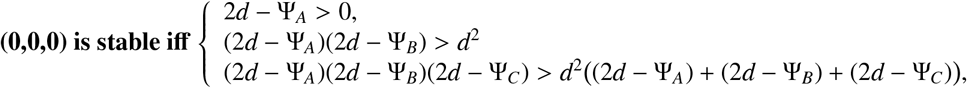

encoding a balance between local stability and diffusion rate. Similarly, for the stability of (1, 1, 1), we introduce: 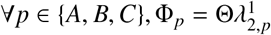. We then reach a condition very similar to the previous one:

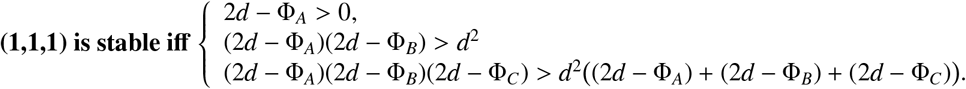

We simulate 3 key scenarios to illustrate system outcomes depending on the above stability criteria: i) (0,0,0) stable and (1,1,1) unstable ii) (0,0,0) stable and (1,1,1) stable hence bistability, and iii) (0,0,0) unstable and (1,1,1) unstable, leading to an interior coexistence point, illustrated in Figure 7A, B, C respectively.

**Figure 7:**
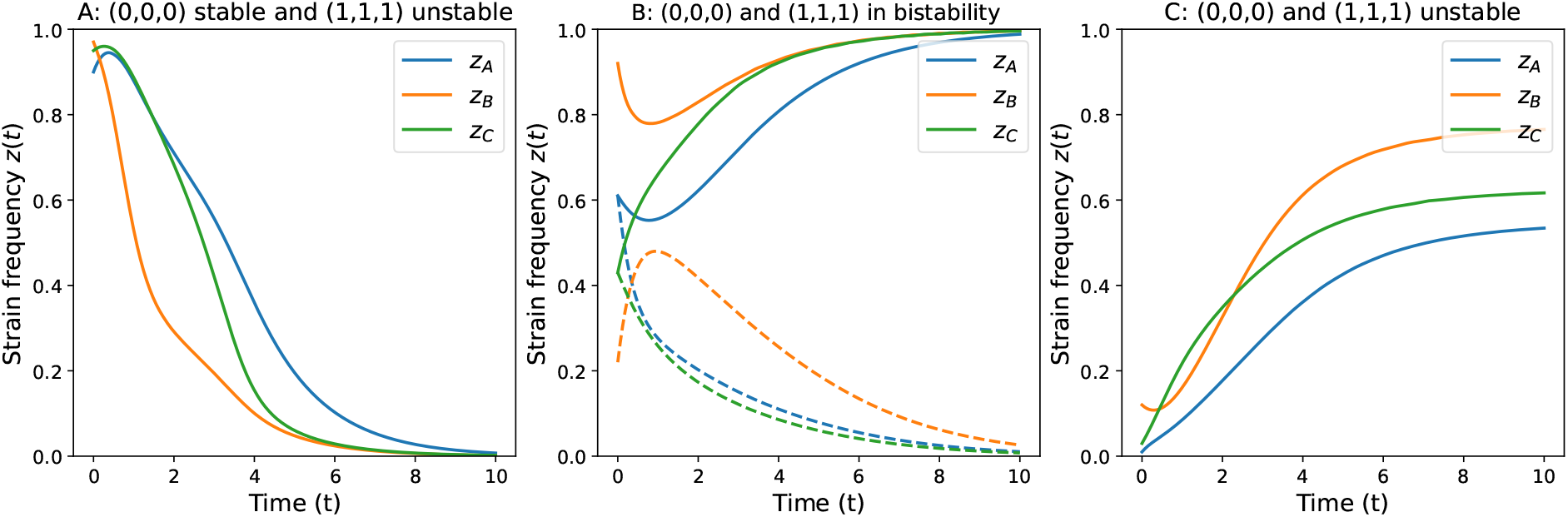
Examples of 2 strain dynamics across 3 patches. **A**. (0, 0, 0) stable and (1, 1, 1) unstable. Dynamics with (*z*_*A*_(0), *z*_*B*_(0), *z*_*C*_(0)) = (0.9, 0.97, 0.95) **B**. Bistability of the marginal equilibria. Solid lines: Dynamics with (*z*_*A*_(0), *z*_*B*_(0), *z*_*C*_(0)) = (0.61, 0.92, 0.43). Dashed lines: Dynamics with(*z*_*A*_(0), *z*_*B*_(0), *z*_*C*_(0)) = (0.61, 0.22, 0.43) **C**. (0,0,0) and (1,1,1) unstable. Dynamics with (*z*_*A*_(0), *z*_*B*_(0), *z*_*C*_(0)) = (0.01, 0.12, 0.03). The system tends to a coexistence between the 2 strains in all three patches. Basic parameter values, equal among patches: β = 5,γ = 0.5, *k* = 0.1,*r* = 0.6, *d* = 1, Θ = 0.084, *R*_0_ = 4.54. Pairwise invasion fitness between two strains are patch-specific: 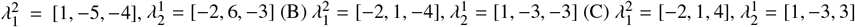.

Overall the simple numerical and analytical illustrations above for low-dimensional cases of the *N*-strain *P*− patch SIS model with co-colonization, serve to highlight the potential for rich nonlinear dynamics of such spatially-extended ecosystems, accessible via spatially modified replicator dynamics (Eq. 6) - where environmental heterogeneity together with local selective forces and host movement combine to generate final competitive outcomes between co-circulating strains. Ultimately space and movement, may favour or hinder coexistence, the answer is crucially: it depends, - and the study of this diversity entangled with environmental gradients should get more complex and fascinating for high number of strains.

## 5. Discussion

Metacommunity ecology is increasingly recognizing and calling attention on the importance of local and regional dynamics for community outcomes (Holyoak et al., 2005; Leibold et al., 2004; Leibold and Chase, 2018). Despite research efforts, the understanding of how different spatial scales interplay to shape metacommunities remains limited. Furthermore the inclusion of environmental heterogeneity into a unified theory of metapopulation dynamics, and the generality of modeled interactions beyond competition (Leibold et al., 2004) are two key areas calling for a long time for mathematical analytical developments, beyond simulation (Thompson et al., 2020).

In this work, we have addressed this challenge, building an *N* −strain *P* −patch metacommunity model, based on SIS dynamics with co-colonization within patches and host migration between patches, incorporating patch heterogeneity and several dimensions of strain variability and interactions. Our model is fully-deterministic, and exploits the methods of time-scale separation under strain similarity (Madec and Gjini, 2020; Le et al., 2023; Le and Madec, 2023). Although inspired by epidemiology, it is easily applicable to other endemic multi-species co-colonization scenarios, developing over a network of interconnected habitats.

Compared to classical multi-patch models, our novelties here are at least two: i) while classical multi-patch models typically adopt the assumption of identical patches, often with symmetric dispersal between them (Levins, 1969; Hanski and Gilpin, 1991; Hanski, 1994; Loreau et al., 2003; Cantrell and Cosner, 2004), with few exceptions, e.g. in heterogeneous networks (Colizza et al., 2007), here the incorporation of explicit patch heterogeneity along several dimensions, together with variable connectivity, makes the model substantially more general, ii) while most multi-patch models incorporate ecological diversity via a few interacting species (Dockery et al., 1998; Okubo et al., 2001; Cantrell and Cosner, 2004; Lerch et al., 2023) (e.g. 2 species predator-prey models) here we encapsulate the full *N* −strain interacting system, in the spirit of adaptive dynamics and frequency-dependent selection with environmental feedbacks Dieckmann and Ferrière (2004).

We studied the model in the case of of low diffusion, where global population structure endemic variables stabilize locally in each patch, on a fast timescale (susceptibles *S*, singly-colonized *I* and co-colonized *D* host prevalences), and a replicator-like equation emerges for local strain frequencies in the slow timescale. The ODE model presented was based on discretization of the PDE framework, originally derived by Le and Madec (2023), but it can also be obtained from first principles and re-application of the slow-fast approach on the multi-patch multi-strain SIS system (Madec and Gjini, 2025).

The power of this approach yields a formulation where patch heterogeneity, in the form of a relevant epidemiological gradient, elegantly appears in the *master* equation (Eq. 6) as a net effective modulation of the movement term in the strain frequency ODE. While the spatial replicator equation regulates local strain frequencies within each patch, it also ‘feels’ global processes operating at larger spatial scales, namely across all the patches in the network, where different prevalence of colonization or co-colonization can magnify or reduce effective strain fluxes across the metacommunity of hosts.

Although we presented the general model formulation for any number of strains and patches, we illustrated the analytical tractability of this model through focusing on the *N* = 2 strain special case. In such case, the system is strongly monotonous, which considerably simplifies the study of the dynamics (Box 1, Section 3.3). For a network of two or three patches, analyzing the original system with population structure and co-colonization becomes prohibitively complex, whereas the replicator equations that emerge after model reduction are strikingly simple. Yet, despite such apparent simplicity, these cases are already able to generate very interesting and non-trivial dynamics, depending on local patch selection outcomes between 2 strains, connectivity, and heterogeneity parameters.

While metacommunity frameworks for *N* = 2 typically focus on the situation of local competitive exclusion (Leibold and Chase, 2018), here we have covered all possible local outcomes, from same-strain competitive exclusion, to opposite-strain competitive exclusion, both-patch coexistence, and bistability. The details of local selection become critical for global outcome especially if the connection is between two opposite-outcome patches or two bistable patches. Indeed, the final outcome depends in nonlinear ways on the balance between magnitude of local outcome stability (Λ_*p*_), speed of local dynamics (Θ_*p*_) and its comparison with movement rate and its net modulation from other patch heterogeneity (net *M* matrix) - an interplay that gets essentially reinforced when the number of strains *N*, is higher, and consequently potential multi-strain outcomes increase hugely in number and complexity, as shown in (Gjini and Madec, 2021).

### Limitations and future directions

Our multi-patch network does not have an explicit notion of distance between patches or population size of patches embedded in it, it only assumes given constant (possibly asymmetric) connectivity rates between all patches in the system. Thus one does not immediately see there the classical gravity models for movement fluxes (Zipf, 1946; Erlander and Stewart, 1990). Yet, extensions of such kind and other special forms could quite flexibly be implemented for specific model applications (see also (Madec and Gjini, 2025) for regularization conditions), making *d*_*pk*_ between any two patches, for example, an adjustable function of their respective sizes and inter-patch distance.

Our model is deterministic, this means it cannot capture how demographic stochasticity can impact metacommunity dynamics (Lerch et al., 2023; Capitán et al., 2017). Next model extensions could account explicitly for different population size between patches and investigate in detail the role of host stochasticity for multi-strain selection dynamics both locally and regionally. We expect in the limit of large population sizes, the model behavior will be similar to the trends and phenomena captured here, but we expect also major differences in small population scenarios, especially when founder-effects are operating at each patch (bistable-bistable connection), and in general a shrinking of coexistence regions in parameter regimes predicted by the deterministic model (Neuhauser and Pacala, 1999).

We also studied local pairwise invasion fitnesses between strains, within a simplified paradigm, namely as decoupled from the global mean-field parameters. However, if as in the original derivation by Madec and Gjini (2020); Le et al. (2023), the invasion fitnesses are mechanistic model-based, and derived from the epidemiological traits between strains, the feedbacks between patch heterogeneity and strain selection could be different. In particular, we expect selection outcomes to be more constrained by such mechanistic couplings, where the λ’s are not entirely free to vary. We have explored some of these scenarios, assuming the invasion fitnesses depend on mean-field parameters of the system, as outlined in Equation 5, in Appendix E. What is fixed is the relative variation between strains. Indeed, if the pairwise invasion fitnesses between strains are trait-based, tracking selective trajectories of specific traits (e.g. mean β, γ, *k*) in a multi-patch model would become possible in the present framework. Because strain frequency evolution is explicitly governed by a replicator-like equation, we can link such selection dynamics directly to the evolution of certain traits locally in each patch and ultimately in the metacommunity; one of which could also be system resistance to invasion.

Although the main goal of this paper is to provide a theoretical underpinning for multi-strain dynamics in a metacommunity of hosts in different patches, our models make also testable predictions in the form of bifurcation diagrams and critical shifts with certain parameters (e.g. Fig.3. These predictions could be tested in laboratory microcosms or with epidemiological field data from co-colonization or co-infection systems with many strains. Indeed, a key challenge in the field of community ecology remains to understand to what extent observational data is informative and how it should be used to infer the underlying community assembly processes (Ovaskainen et al., 2019).

Examples of biological systems for applications and analogies could be systems with SIS-type (or logistic growth) dynamics at their core (Levins, 1969; Gyllenberg et al., 1997), co-colonization processes and diversity of co-circulating variants across different sites, for example endemic dynamics of antibiotic-resistant and antibiotic-sensitive strains between two interconnected habitats, dynamics of two virus variants across two body compartments, pneumococcus serotype dynamics across geographic locations, co-colonizing species dynamics across two locations differing in temperature, humidity or other physical parameters, and even cancer metastatic dynamics viewed under the lens of metapopulations (Halkola et al., 2022). In such contexts, the questions of habitat fragmentation and biodiversity, climate change and biodiversity, antibiotic pressure gradient and selection in metapopulations could be explored further. The role of movement, or dispersal for biodiversity could be studied for specific cases, and the model framework provided here could help reconcile conflicting patterns, namely that sometimes movement could help promote biodiversity at the regional scale and other times it could hinder it. We believe mechanisms of global vs. local coexistence, captured here, can be put in practice for effective management of biodiversity and ecosystem health in a wide variety field of applications in the future, from agriculture to biomedicine.

## Acknowledgement

We thank Thi Minh Thao Le for helpful discussions at the beginning of this work. The study was supported by the Portuguese Foundation for Science and Technology (FCT grant number 2022.03060.PTDC), by CEMAT (fellowship grant under UIDP/04621/2020 to M.M.) and by the ERASMUS EU programme (fellowship to L.P).

## Funding statement

The study was supported by the Portuguese Foundation for Science and Technology (FCT grant number 2022.03060.PTDC), by CEMAT (fellowship grant under UIDP/04621/2020 to M.M.) and by the ERASMUS+ EU programme (fellowship to L.P).

## Appendix A. Patch heterogeneity in mean-field parameters and ν_*pk*_

### Explicit parameter values: Θ, ν for each patch

In the 2-patch case, assuming without loss of generality that demographic turnover is equal in both patches (*m*) and the mean-field environmental difference between patches is given by basic reproduction number variation and single - to coinfection ratio variation:

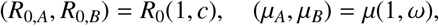

in the system above the speed parameters can be expressed as:

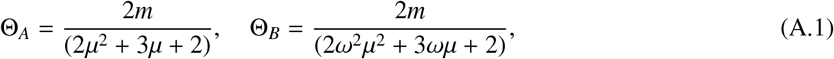

and the global migration contribution parameters *nu*_*AB*_ and *nu*_*BA*_ are given by:

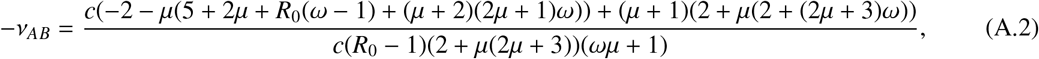

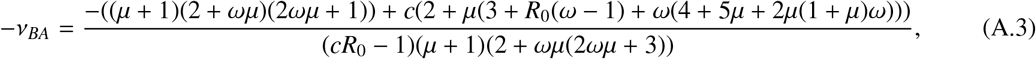

which clarifies their dependence on global mean-field parameters in each patch and their ratios, but defies a straight-forward immediate understanding or prediction of effects. That is why we resort to simulations.

## Appendix B. Examples for 2 patch system for *N* = 2

### Case of anti-symmetric invasion fitnesses in each patch

Let’s consider a simple case of:

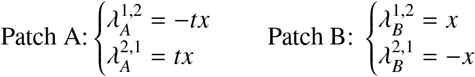

with *t, x* > 0 so that we keep the necessary fitness structure for opposite exclusion. We can control the stability difference of opposite strain exclusion in each patch by varying the coefficient *t*. We assume patches have equal ratios of single to co-infection µ (ω = 1).

### The case c = 1: Patches have equal R_0_

Assuming equal patches in both *R*_0_ and µ parameters: *c* = 1, ω = 1, we have the following. This becomes a very simple system of equations and can be completely described analytically. Besides the trivial global exclusion equilibria, there exists one feasible coexistence equilibrium

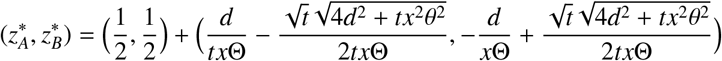

which can be understood as a perturbation from the average point 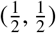 and if *t* = 1 then 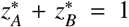. The conditions for existence and stability of this point are identical and can be described as:

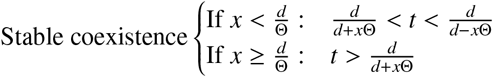

So for fixed patch-specific parameters, there is a critical value of migration between patches *d* above which we go from a system of 2 exclusion equilibria (0,0) and (1,1), to a system with 3 equilibria including coexistence in both patches. This critical value is given by: 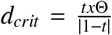. And for the marginal equilibria, where the same strain gets fixed in both patches, we have:

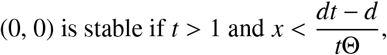

and

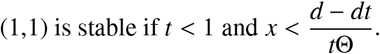

The conditions can be interpreted as comparing the relative strengths of stability of each exclusion equilibrium in each singular patch when uncoupled and identifying a critical lower bound for migration rate between patches.

(0, 0) is the equilibrium where Patch A trend dominates ultimately in both patches in the coupled system, and is achieved for relatively higher stability of the *z* = 0 equilibrium in A vs. the *z* = 1 equilibrium in B, namely *t* > 1. This, together with a sufficiently high diffusion rate 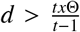 guarantees strain 1 loses in both patches, and strain 2 being the sole strain that persists in the coupled system. Similar arguments apply to the (1, 1) equilibrium: it is achieved for *t* < 1 and again its stability requires 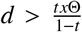, These conditions, taken together indicate that strong diffusion between two originally opposite exclusion patches (sink -source) favours exclusion, eventually one or the other strain will be the sole winner in both patches.

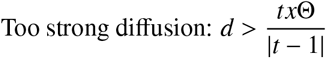

Conversely, for intermediate rate of diffusion between such patches, coexistence of both strains in both patches is possible and stable:

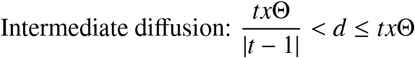

In such coexistence, there can be biases in relative abundances, driven mainly by the asymmetric stabilities of each original exclusion in the respective patches under no coupling (*t* -:t 1).

The above conditions imply that bistability between the marginal equilibria (0, 0) and (1, 1) is impossible under these assumptions. Furthermore, once there is coexistence, the marginal equilibria become unstable so there is also no bistability between a coexistence and an exclusion equilibrium. This means that the entire qualitative behaviour of the system can be completely described by the verification of the previous three conditions relating *t, x, d* and Θ.

### The case c > 1: Patch B has higher R_0_

When *c* > 1, hence when patch B has a bigger *R*_0_ than patch A, we are still able to find closed form solutions for the coexistence equilibrium, but it is no longer straightforward to obtain the corresponding conditions for stability. The complications come from ν_*AB*_ -:t 0 and ν_*BA*_ -:t 0 anymore. Similarly, as before, this equilibrium is given as a perturbation from 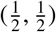 by

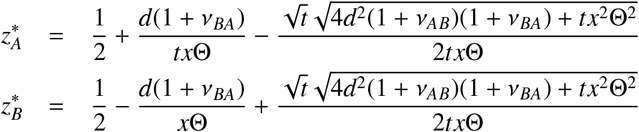

and it should always exist. Because the direct conditions for its stability are not immediately tractable analytically, we can resort to reasoning about the stability of the marginal equilibria. Here, we have:

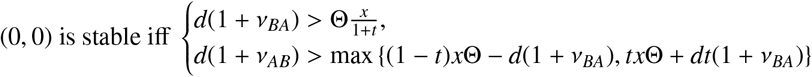

and

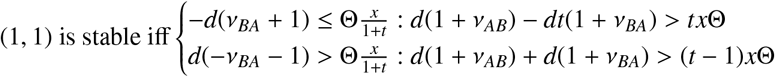

And so the outcome of the system might be predicted from these two conditions. These criteria demonstrate that the ultimate fate of the E1-E2 system will be decided by an interplay of all parameters together, including the migration rate *d*, the difference between patches in global mean-field epidemiological parameters manifested as ν_*AB*_ and ν_*BA*_ but also from the exact magnitudes of stability of exclusion in each patch, namely the parameters *x* and *t*. Notice also that even when the two patches are equal in some parameter, here µ, and as a consequence in Θ, this parameter, may still have an effect on global outcome; in this case Θ represents the speed of local dynamics and it interacts with *d* the speed of global mixing of the two strains in the system.

## Appendix C. The 2-strain 3-patch system with homogeneous patches

The analysis performed in this special case requires the following additional hypothesis: All 3 patches have the same neutral model. In other terms, all patches are identical for fast timescales : β_*A*_ = β_*B*_ = β_*C*_, γ_*A*_ = γ_*B*_ = γ_*C*_… Only pairwise invasion fitnesses and slow variables *z* are now patch patch-specific. This framework enables in particular to take out the advection terms ν_*p*_ = 0.

In this context, the equation for the 3-patch system is:

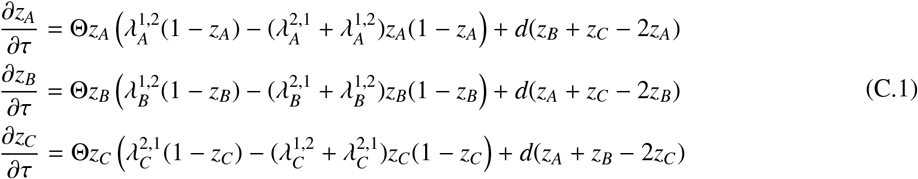

We now propose to study the stability of the two trivial equilibria of this system. Our two Jacobians are written on these points:

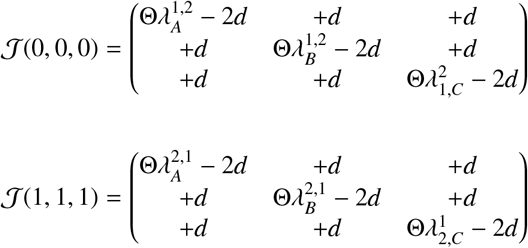

We get two symmetric matrixcs. And this is where the additional hypothesis made in the beginning of the section makes sense (without it, the Jacobians would not be symmetric).

Now, we want to see, under which conditions, each of these two Jacobians has strictly negative eigenvalues, which is equivalent to them being symmetric definite negative. Noticing this enables us to use an algebraic criterion, which is going to simplify a lot the study of stability : Sylvester’s criteria.

### Sylvester’s criteria

*Sylvester’s criteria* : “Let A = (*a*_*ij*_)_1≤*i,j*≤*n*_ be a real symmetric matrix. Then, A is positive definite if and only if the determinants of the *n* leading principal submatrices A_*p*_ = (*a*_*ij*_)_1≤*i,j*≤*p*_ for *p* = 1, …, *n* are all strictly positive, that is, if all the leading principal minors of A are strictly positive.”

*Proof*. We will work in a vector space of dimension n E. Since A is a real symmetric matrix, it is diagonalizable with transition matrix Q. Let us introduce its real eigenvalues (λ)_*i,i*∈1,…*n*_, and their eigenvectors (*e*_*i*_)_*i*∈1,…*n*_. This means we have A = Q^*T*^ ≱Q, where ≱ = *Diag*(λ_1_, λ_2_, …λ_*n*_).

For each *e*_*i*_, let us say E_*i*_ = *Vect*(*e*_*i*_). Since A is diagonalisable one can write:

E = E_1_ ⊕ E_2_ ⊕ E_*n*_

A positive definite enables us to write: ∀*i* ∈ ⟦ 1, *n* ⟧, λ_*i*_ > 0.

Now we are going to consider the reductions of endomorphism A on the vector spaces E_1_, E_1_ ⊕E_2_,…E_1_ ⊕E_2_⊕ E_*n*_.

Each of them will guarantee the positivity of the determinant of a leading submatrix.

For instance let us look at A_*p*_ = (*a*_*ij*_)_1≤*i,j*≤*p*_, for *p* ∈ ⟦1, *n*⟧ . In order to prove that its determinant is > 0, we have to look at the reduction of endomorphism A on E_1_ ⊕E_2_ ⊕ …E_*p*_

Then we have that : *det*(A_*p*_) = *det*(Q_*p*_)^2^ ∏_*i*∈_⟦_1,*n*_⟧ λ_*i*_ > 0, where Q_*p*_ = (*q*_*ij*_)_1≤*i,j*≤*p*_ and the result is proven.

By considering the same reductions of endomorphism A, we get that λ_1_ > 0, λ_1_λ_2_ > 0, … Π_*i*∈ ⟦1,*n*_⟧ λ_*i*_ > 0.

This leads to: ∀*i* ∈ ⟦1, *n*⟧, λ_*i*_ > 0, which leads to A definitive positive.

### Stability of (0,0,0)

We are going to use this result in the following way : an equilibrium is stable if and only if all the leading principal minors of the opposite Jacobian matrix at this point are strictly positive.

Here is our Jacobian:

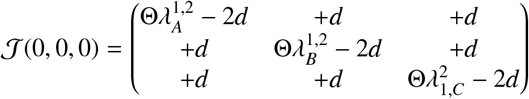

Its opposite is written:

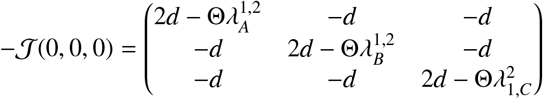

The criteria guarantees (0,0,0) to be stable if and only if:

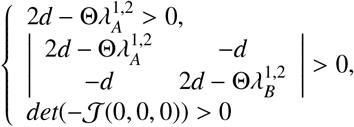

Now let us introduce:

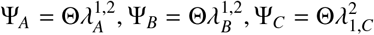

We have: **(0**,**0**,**0) is stable for the network i**ff

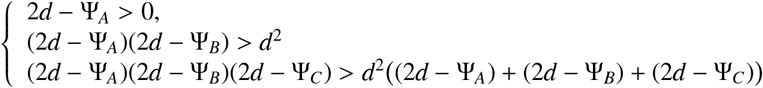

Now the following table predicting the outcome of a 2-strain scenario allows to make a relevant remark

| Pairwise invasion $(\lambda_1^2, \lambda_2^1)$ | Stable positions | Strain frequencies at eq. |
| --- | --- | --- |
| (+, +) | Coexistence equilibrium | $z_1^* = \frac{\lambda_1^2}{\lambda_1^2 + \lambda_2^1}, z_2^* = \frac{\lambda_2^1}{\lambda_1^2 + \lambda_2^1}$ |
| (+, -) | (1) | $z_1^* = 1, z_2^* = 0$ |
| (-, +) | (0) | $z_1^* = 0, z_2^* = 1$ |
| (-, -) | (0) and (1), bistability | $z_1^* = 0$ or $z_2^* = 1$ |

### Corollary

In particular if in all three isolated patches (0) is stable then 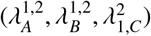 is in 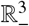, then (Ψ_*A*_, Ψ_*B*_, Ψ_*C*_) is in 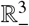, then all three conditions are verified, which implies that (0,0,0) is stable.

In short our triple criteria enables to show the intuitive: ‘(0) stable in every isolated patch’ ⇒ ‘(0,0,0) is stable for the network’

### Remarks

- It is worth mentionning that the three patches are interchangeable, meaning that their labels A, B and C can be taken arbitrarily.
- Besides, it can be interesting to see the impact of network connectivity on the “stability volume”.
  − **Strongly connected networks**: If *d*→ + ∞ then the volume defined tends to the semi-space x+y+z¡0. (0,0,0) will be stable iff

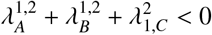

that is iff (0) is “stable on average”
  − **Weakly connected networks**: If *d* → 0 then the volume defined tends to R^3^ . At the limit the patches aren’t connected and so (0,0,0) is stable iff (0) is stable for every patch.

### Stability around (1,1,1)

Now the Jacobian studied is::

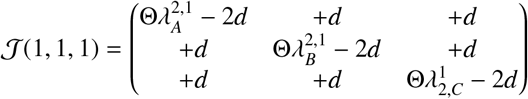

In the same spirit as last section, we write its opposite in:

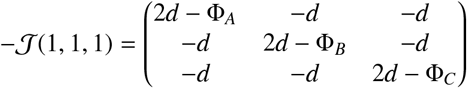

Where we have introduced : ∀ *p* ∈{*A, B, C* }, 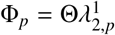

We then reach a condition very similar than the previous one.

**(1**,**1**,**1) is stable for the network i**ff

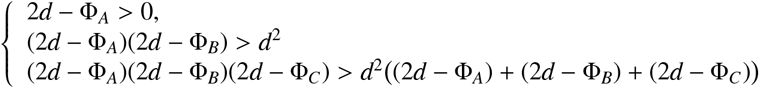

*Corollary*. In particular if in all three isolated patches (0) is stable then 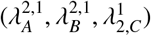 is in ℝ_−_^3^, then (Φ_*A*_,Φ_*B*_,Φ_*C*_) ℝ_−_^3^, then all three conditions are verified, which implies that (1,1,1) is stable for the network. In short our triple criteria enables to show the intuitive: ‘(1) stable in every isolated patch’ ⇒ ‘(1,1,1) is stable for the network’.

## Appendix D. Supplementary figures and further illustrations

Here, we test the accuracy of our predictions in Figure 2 of the paper, by picking points in each region and solving the system numerically (Figure D.1).

**Figure D.1:**
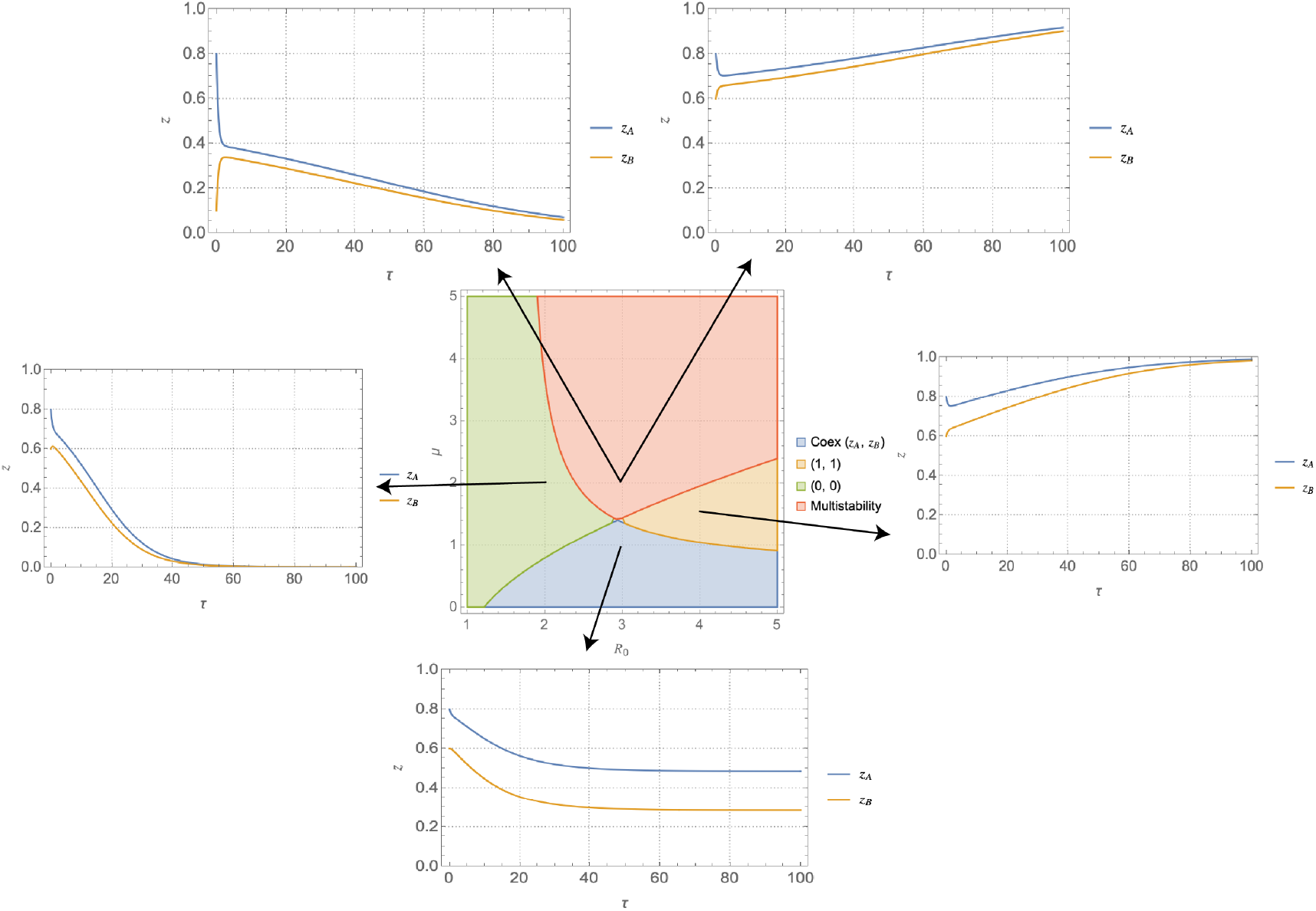
Solutions of the system associated to different point of the (*R*_0_, µ)-plane for parameters as in figure. The regions of bistability can arrive at either exclusion, depending on initial conditions. The Cases of (1, 1) and (0, 0) behave as expected, although the resulting dynamics could be more complicated. The coexistence region always maintains the diversity of strains over both patches, although the relative frequencies of each strain will change for the points inside the region.

Finally, we can illustrate a sample solution for one of these points in the regions where more than one coexistence point exists (Figure D.3), for *R*_0_ = 3.2 and µ = 1.4.

We can study the effect of host movement rate on this particular system by keeping every parameter as before but changing *d* (Figure D.4). In this example, we can see that increasing the magnitude of movement *d* hinders diversity, in this particular system. The regions for coexistence shrink as *d* increases. These regions naturally do not allow us to understand whether the relative frequencies in these coexistence regions become more homogenized as *d* increases but it is clear that higher movement does not, for this set of invasion fitnesses, favour these two strains to coexist.

**Figure D.2:**
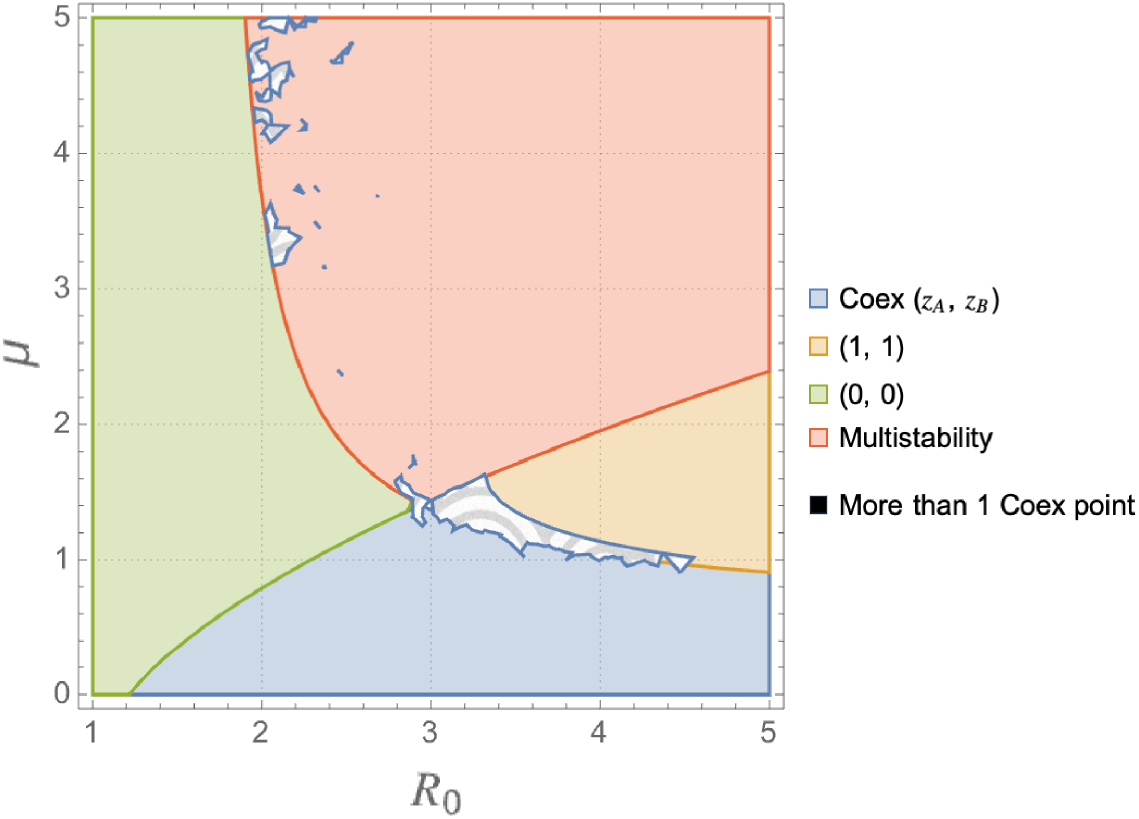
Regions of competitive outcomes for a special system with a test on uniqueness of a coexistence point. In the textured areas, there exists more than one coexistence point, meaning the behavior of the system can be more complicated. It is over these regions that marginal equilibria stability is not sufficient to capture all dynamics.

**Figure D.3:**
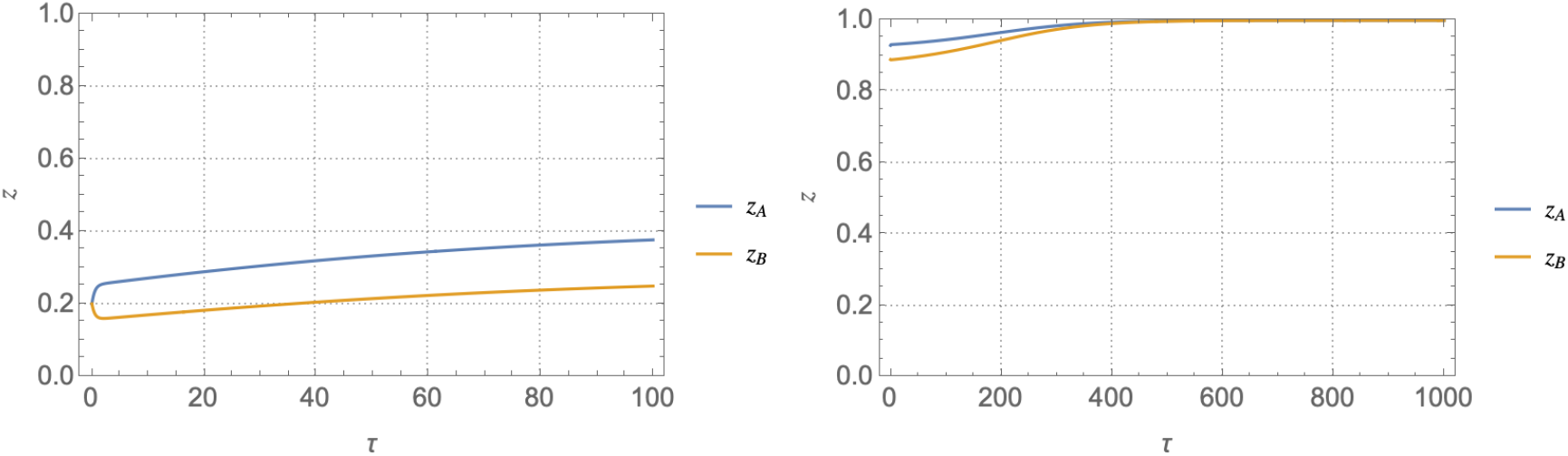
Verification of the complex behavior of the system when there exists more than coexistence point. (1, 1) is stable however there is another stable coexistence point. Depending on initial conditions (left: *z*_*A*_(0) = *z*_*B*_(0) = 0.2, right: *z*_*A*_(0) = 0.93, *z*_*B*_(0) = 0.89), the system will either fixate strain 1 or approach the stable interior coexistence. Strain 2, however, can never be fixated.

**Figure D.4:**
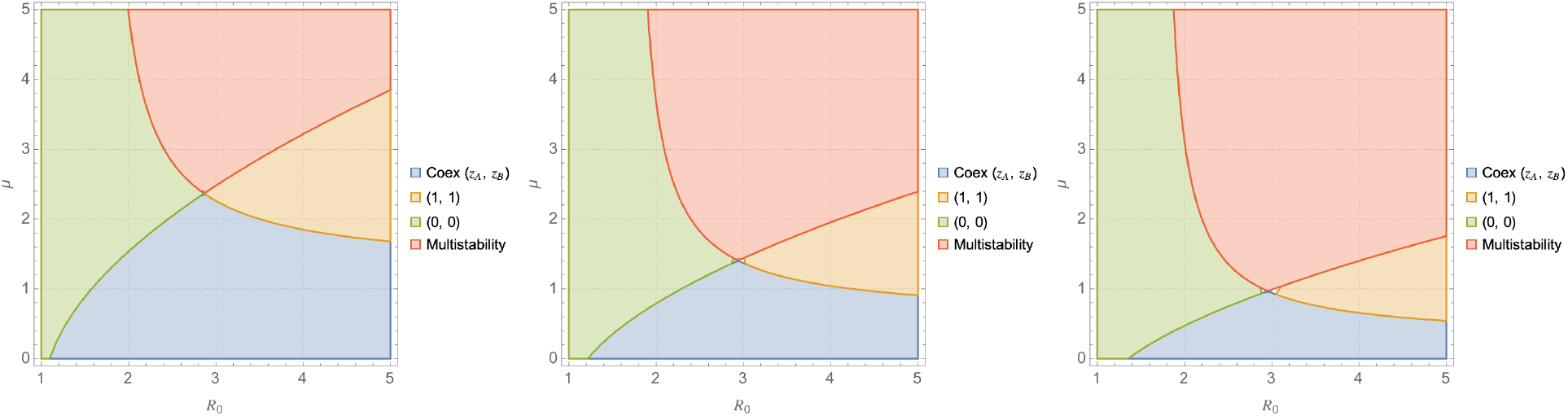
Effect of movement on the system with (0)-(1) patch connection, and parameters as in Figure 2 of paper. Left: *d* = 0.5, middle panel: *d* = 1, right: *d* = 1.5. As host movement increases the region of unique stable coexistence decreases.

## Appendix E. Mechanistic invasion fitnesses and predicting 2-strain outcomes in the (*c*, ω)-plane

In the main bulk of the paper, we have considered the invasion fitness to be 4 free parameters from their specific patch’s parameters. This is, however, not the most realistic definition, when recalling the derivation from the SIS model. Here we will analyze the effect that connecting invasion fitness to the environment has on the resulting dynamics of the system. The mechanistic definition of invasion fitness (Madec and Gjini, 2020), assuming only trait variation in α_*ij*_, is

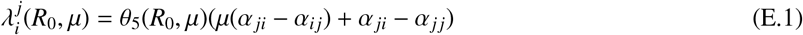

and recall that in the non-spatial case the relative sign of the invasion fitness’ is enough to determine the competitive behavior of the system (Figure E.5).

**Figure E.5:**
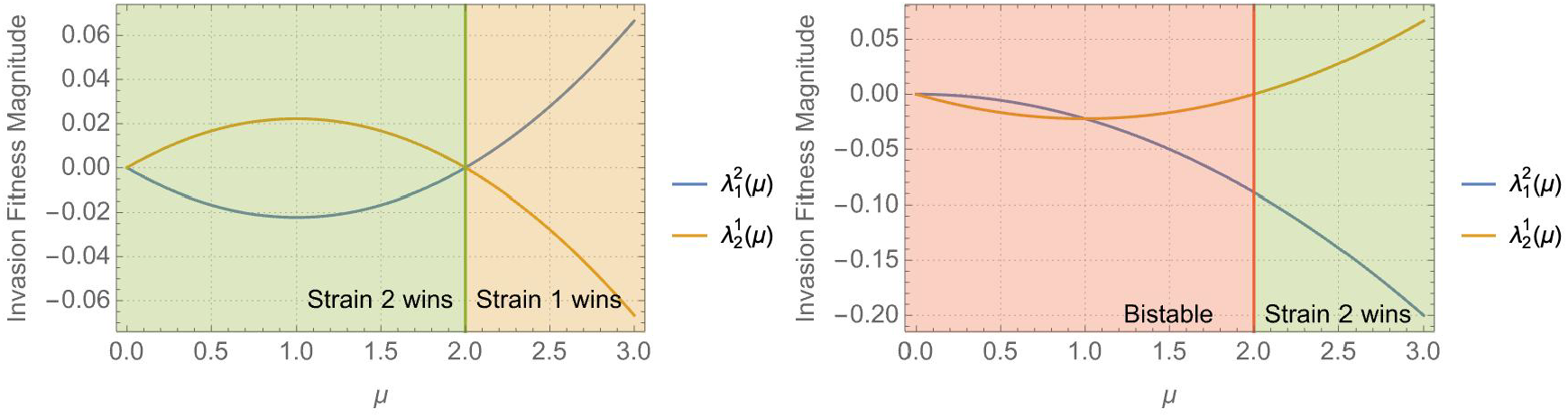
Relation between the competitive outcomes of a single uncoupled patch and the invasion fitness functions of µ, for fixed α_*ij*_’s.

With the usual framework, by fixing µ_*A*_ = µ and varying ω we will be able to shift the system into heterogeneous regimes of competitive preference, that is, the patches will have different competitive outcome preference as we vary µ. This allows us to connect the invasion fitness’ to “environmental” properties of patches, in particular, to the ratio of single-to co-infected prevalcne µ. Of course, we still have to fix the set of α_*ij*_ over both patches, but ultimately this can be done in a systematic manner directed at having different types of dynamics arise from sweeping the values of ω, that is, changing µ_*B*_ in such that patch B changes its separate uncoupled outcome. Recall still that α_*ij*_ in the full epidemiological model 1 is associated to the neutral altered factor of susceptibility *k* between strains and the strain dissimilarity ϵ

**Figure E.6:**
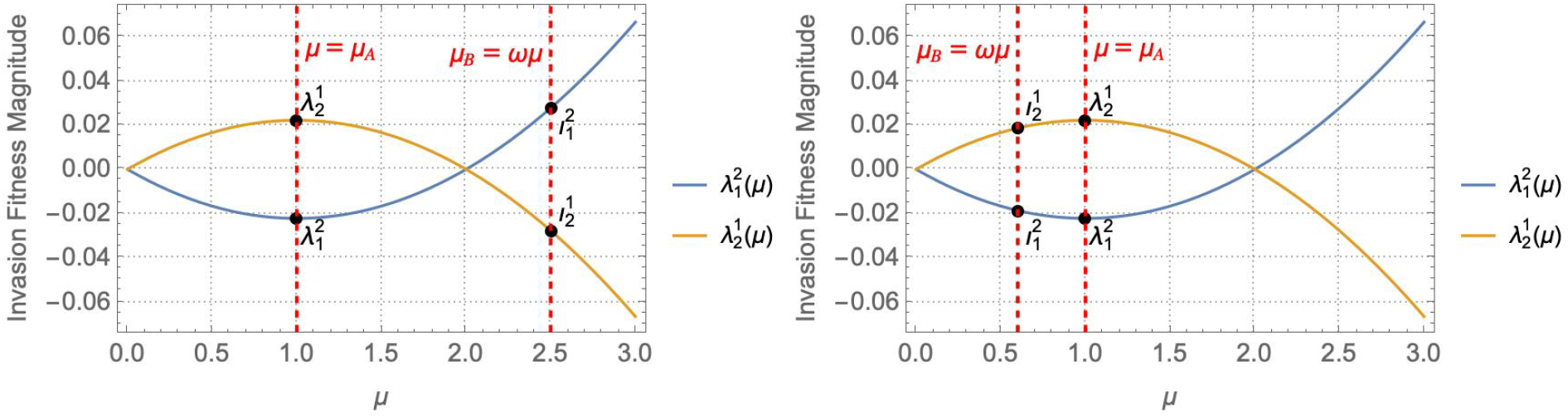
Respective values for mechanistic invasion fitness under full model derivation, in µ and ω framework. Left: ω = 2.5, right: ω = 0.6

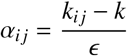

As before, we will assume

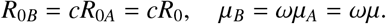

We decided to fix *R*_0_, as it does not appear explicitly in 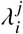, and µ; and let *c* and ω vary. The motivation for this approach comes from the last section: in fixing µ we can keep the parameters of patch A fixed and, by varying ω, we are able to possibly change patch B into a different parameter regime (Figure E.6). Furthermore, the type of selection change happens in a way which is connected to the “environmental” parameters and not taken arbitrarily, as before. That is, for every point in the (*c*, ω)-plane there are a set of corresponding invasion fitness’; whereas in the paper (e.g. Figure 2) the four invasion fitness where fixed and constant over the whole (*R*_0_, µ)-plane.

We split the analysis into various qualitative cases of interest that arise from certain choices of α_11_, α_12_, α_21_ and α_22_. This constraint will restrict the quality of patches we can connect. It is for example, impossible to connect a coexistence patch with a bistable one. Under this framework, not much analysis can be carried out analytically. Instead, we have used `Mathematica` to evaluate the usual Routh-Hurtwiz criteria for the stability of the marginal equilibria for points in the (*c*, ω)-plane. We need to fix *R*_0_, µ, *d*; and we will look into specific choices of α_*ij*_ so as to enable different selection scenarios. We have also taken α_*ij*_’s to be equal between the two patches. Throughout these various cases, we have kept the same values for the fixed parameters. These are resumed in Tables below:

**Table E.7:** Fixed parameters for each patch.

| Table E.5: * |  |  |
| --- | --- | --- |
| Patch A | Fixed Parameter | Value |
| | $R_{0A} = R_0$ | 1.2 |
| | $\mu_A = \mu$ | 1 |
| | $d$ | 0.2 |
| | $m$ | 1.5 |

Table E.7: Fixed parameters for each patch.
| Table E.6: * |  |  |
| --- | --- | --- |
| Patch B | Fixed Parameter | Value |
| | $R_{0B}$ | $1.2c$ ($c$ is varied) |
| | $\mu_B$ | $\omega$ ($\omega$ is varied) |
| | $d$ | 0.2 |
| | $m$ | 1.5 |

### (0) in patch A and patch B transitions from (0) to (1)

Here we have looked at a case where the invasion fitness functions allow the patches to shift **(0), (0)** to **(0), (1)** as a function of µ. Patch A is always of type **(0)** but as we vary ω we force patch B to become of type **(1)** (Figure E.7). For a set of fixed 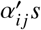

**Figure E.7:**
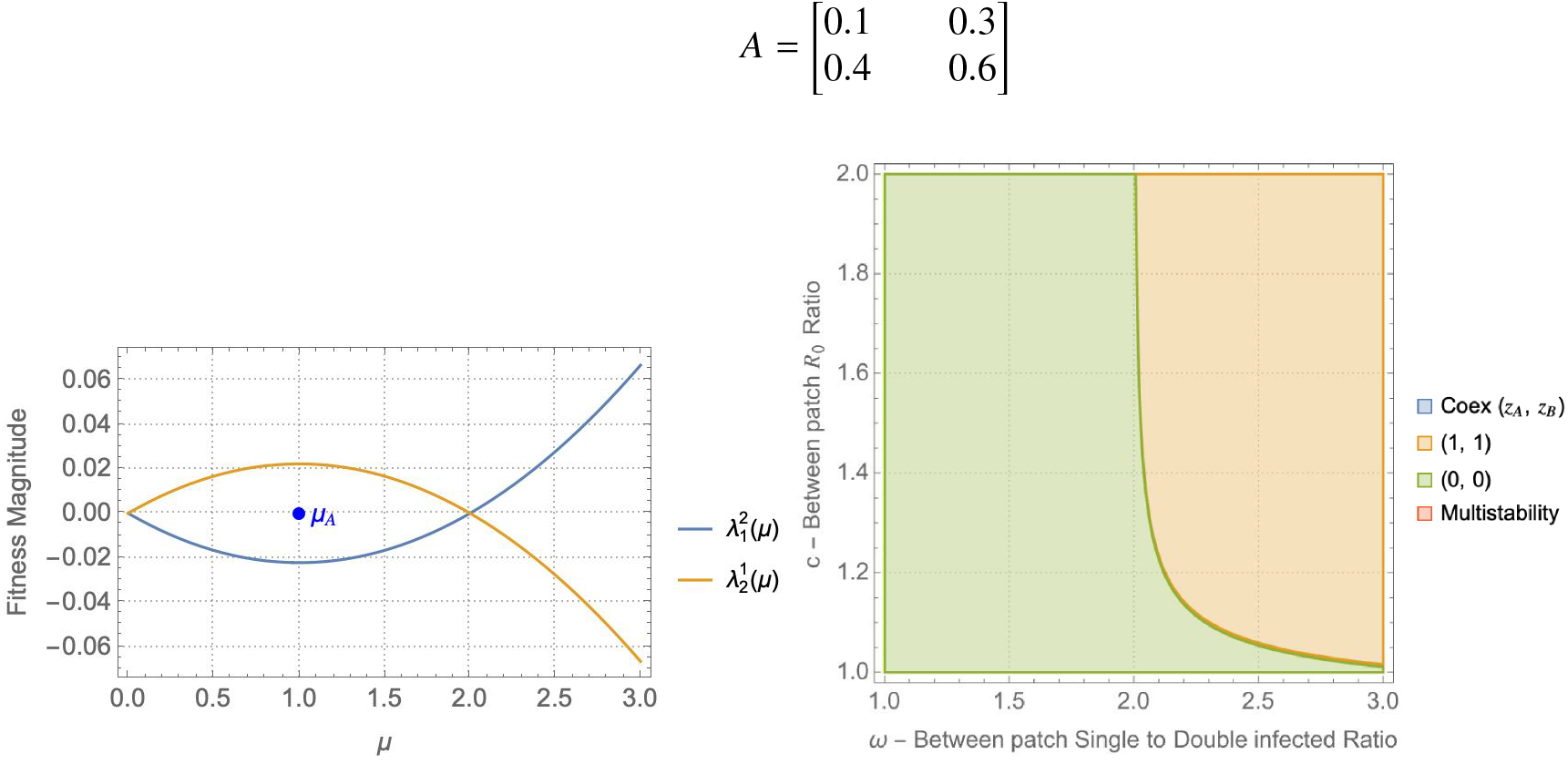
Possible outcomes arising from coinfection susceptibility variation *A*. Left: Invasion fitness curves. Right:Regions of stability in the (*c*, ω)-plane. The system can go from a **(0, 0)** to a **(1, 1)** outcome, meaning that the fixation of strains will depend on the size of ω and *c*.

We can take a closer look at how the transition from **(0, 0)** to **(1, 1)** happens in the (ω, *c*)-plane and see there is actually a small boundary region of coexistence (Figure E.8.)

**Figure E.8:**
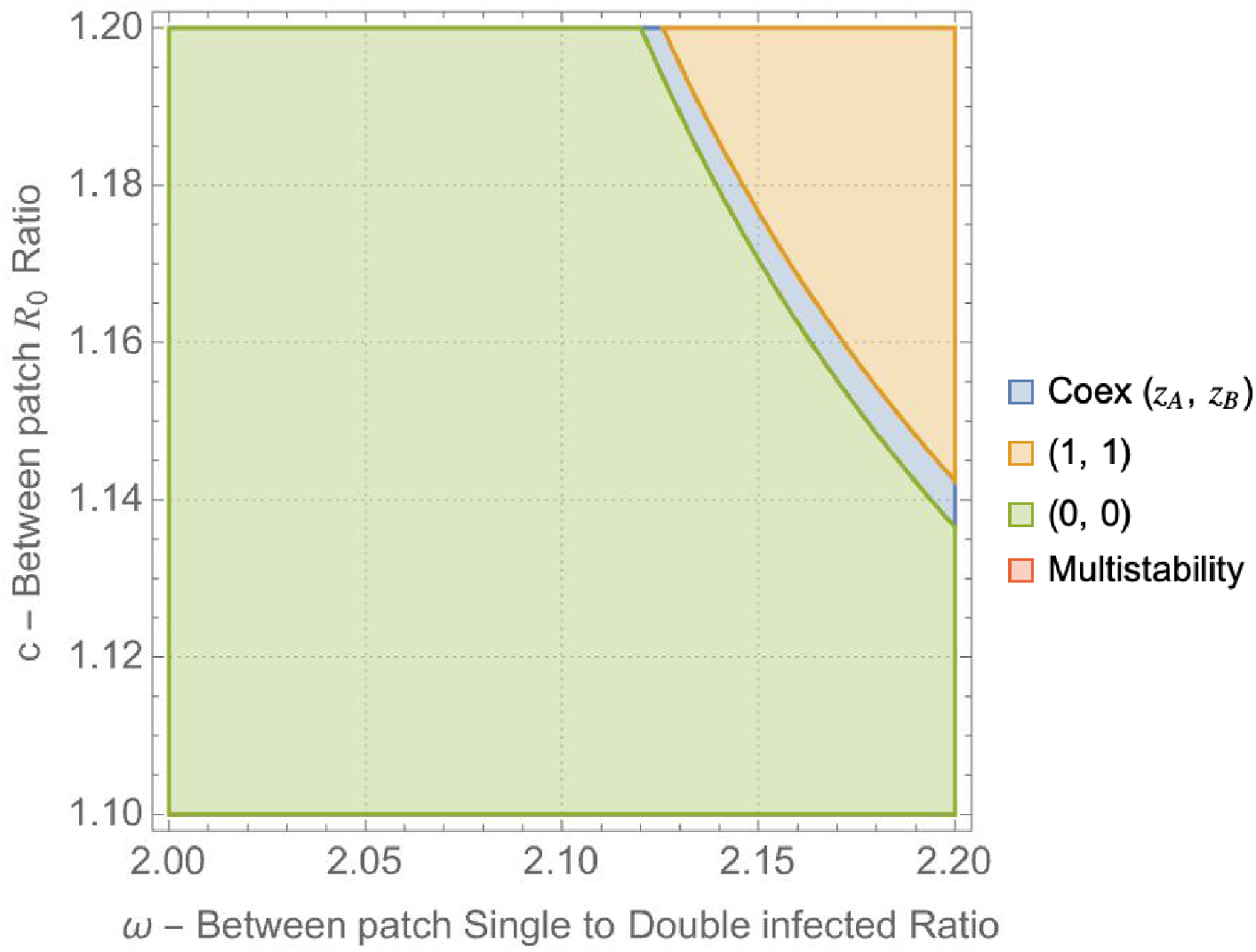
The transition from **(0, 0)** to **(1, 1)** is done via a thin region of coexistence.

### Patch A is multistable and patch B transitions from multistability to (0)

This invasion fitness structure is similar as before, but patches move from a multistable competitive outcome **(0**/**1), (0**/**1)** to a **(0**/**1), (0)** competitive outcome (Figure E.9). Here

**Figure E.9:**
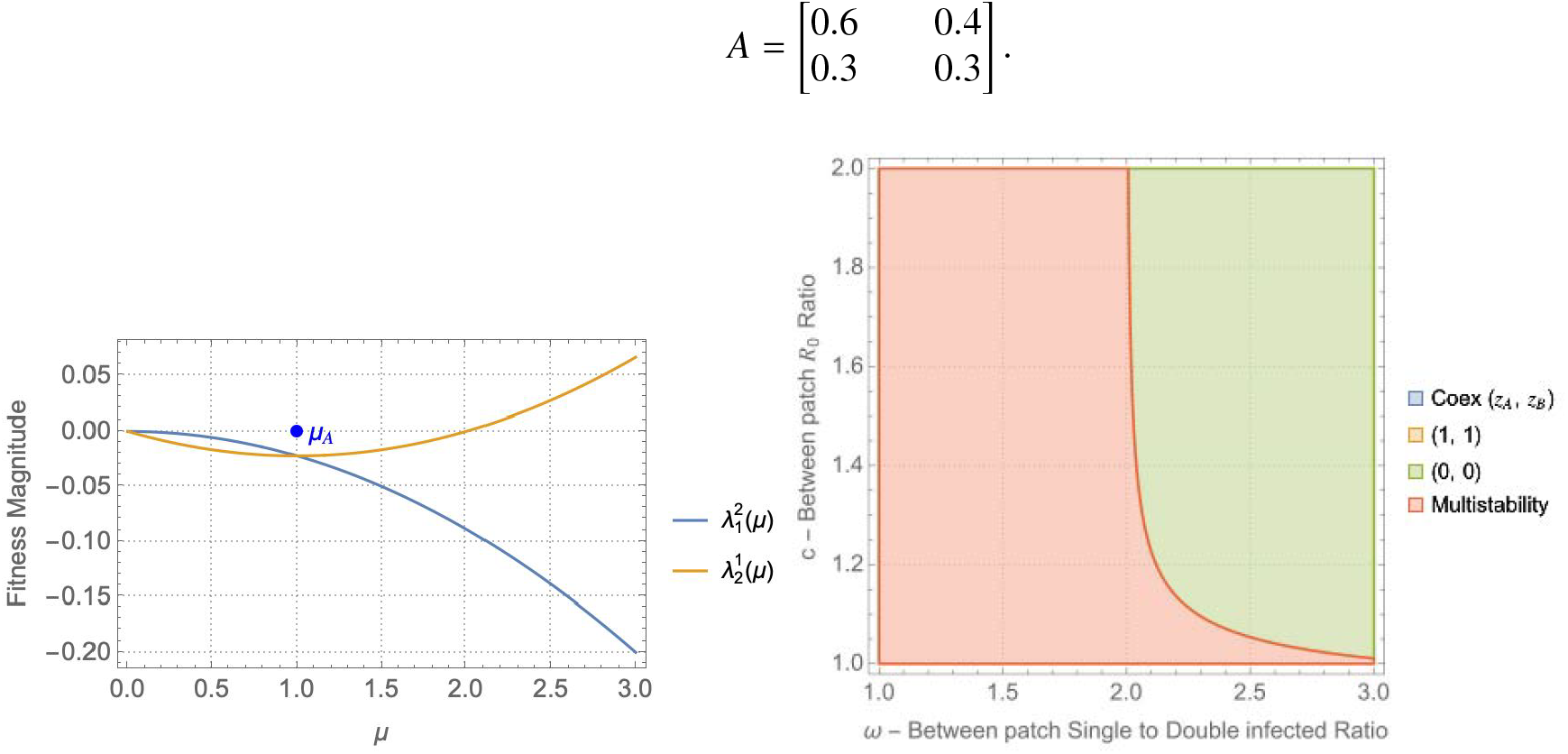
Possible outcomes for the chosen *A*. Left: Invasion fitness curves. Right: Regions of stability in the (*c*, ω)-plane. Both multistability and **0, 0** are possible, depending on ω and *c*.

### There is coexistence in patch A and patch B transitions from coexistence to (1)

Taking the symmetric invasion fitness from the previous case allows to shift the patches from a coexistence coupling **(*z*), (*z***^**′**^) to a **(*z*), (1)** coupling (Figure E.10). That is

**Figure E.10:**
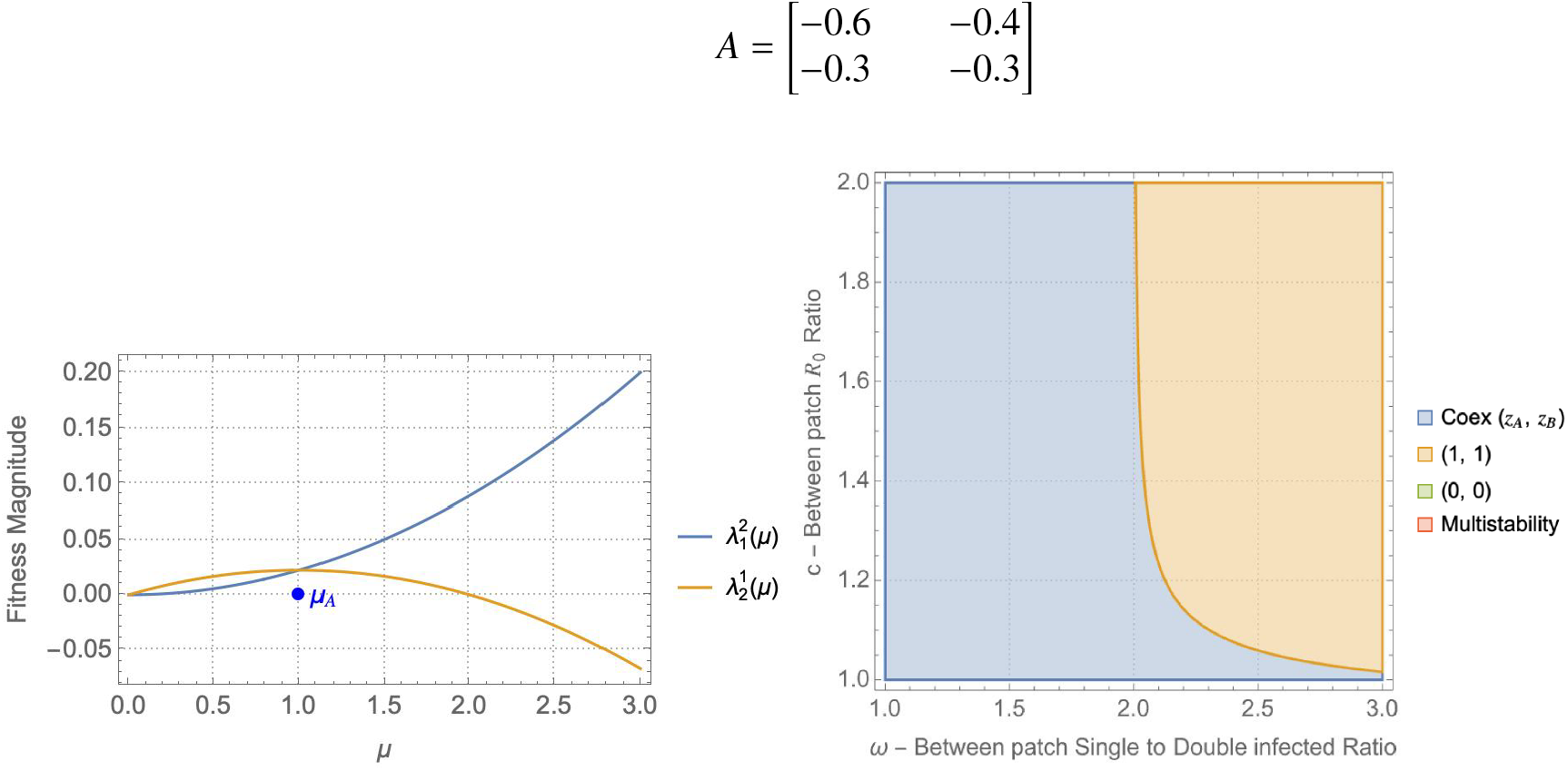
Possible outcomes from fixing *A*. Left: invasion fitness curves. Right. Regions of stability in the (*c*, ω)-plane. We now can arrive at either 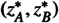or **(1**, 1).

### (0)in patch A and patch B transitions from 3 outcomes

And finally, perhaps more interestingly, we can define invasion fitness function that allow for three different scenarios depending on the choice of ω. Now we have taken

**Figure E.11:**
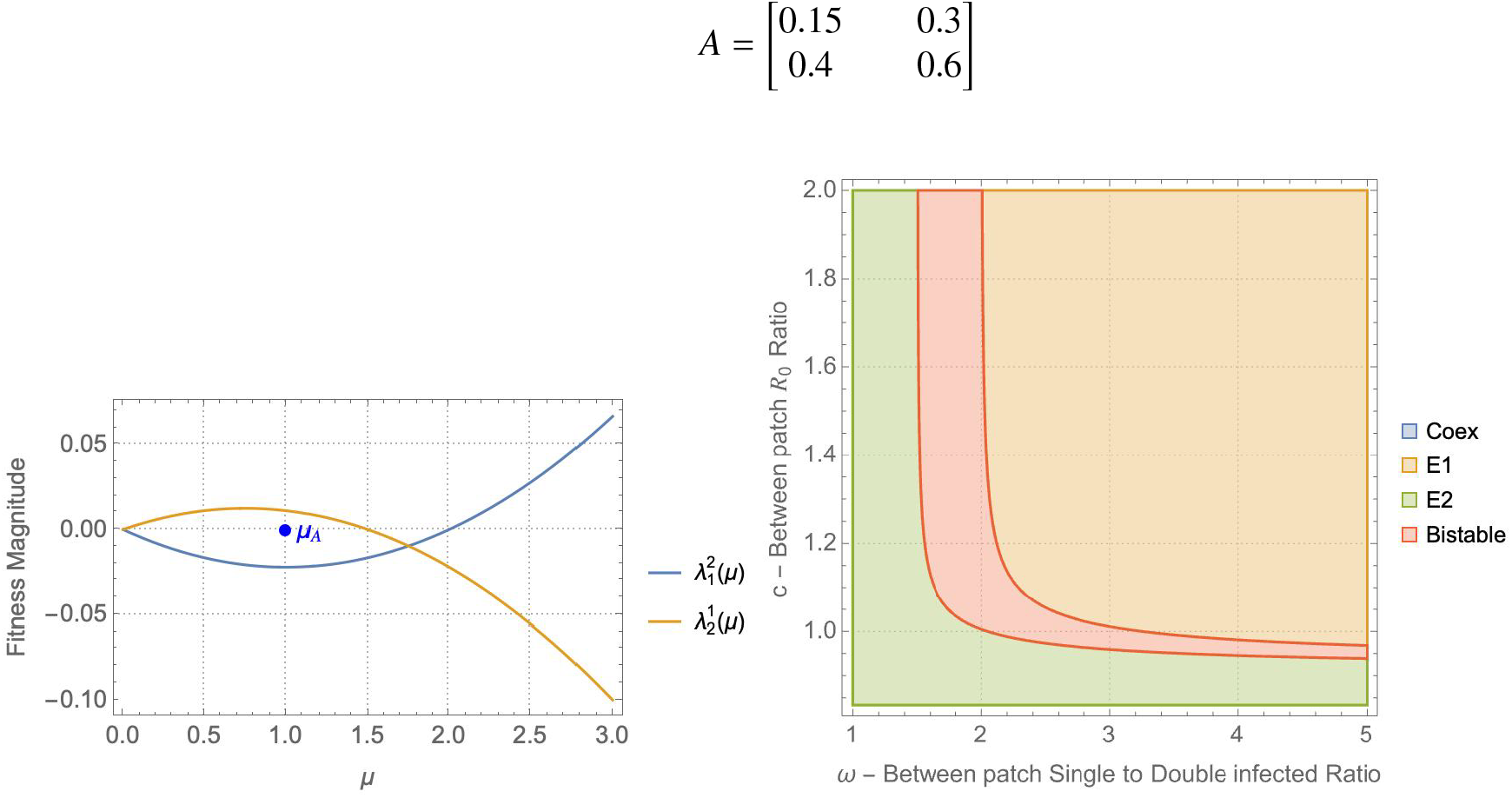
Three possible outcomes from coupling. Left: Invasion fitness curves. Right: Regions of stability in the (*c*, ω)-plane. Depending on ω and *c* we can arrive at either **(0**, 0), multistability or **(1**, 1). Increasing *c* and ω favors the fixation of the first strain. Between the two mono-exclusions there is a region of multistability.

There are clearly 3 regions for µ, associated to three different types of coupling in Figure E.11.

1. µ ∈]0, 1.5[ we have **(0), (0)** coupling;
2. µ ∈]1.5, 2[ we have **(0), (0**/**1)** coupling;
3. µ ∈]2, +∞[ we have **(0), (1)** coupling;

Depending on which of these regions of µ we are placed in the (ω, *c*)-plane, the effect of *c* will be significant or not in the global coupled outcome. To conclude, the mechanistic connection of invasion fitness to environmental parameters as in the strict model derivation (Madec and Gjini, 2020) reduces the possible outcomes of the system. Primarily

- Not all types of patch couplings can be achieved, that is, we can only connect certain types of patches whose invasion fitness functions shift from one competitive regime to another and functions do not allow for all possible regimes to be connected via µ.;
- For the possible couplings, variation in *c* and ω is enough to obtain different competitive outcomes
- The maximum number of possible outcomes found for a given system was 3.
- The 5-dimensional trait variation governing Λ as derived in (Le et al., 2023) is interesting to be studied in the future.

## Notes

### Competing Interest Statement

The authors have declared no competing interest.

### Summary of Updates

Added graphical diagram of the model Clarified theoretical results for special cases Modified slightly the introduction and discussion Rephrased and clarified several points

## References

Alizon, S., 2013. Co-infection and super-infection models in evolutionary epidemiology. Interface focus 3 (6), 20130031.

Arditi, R., Lobry, C., Sari, T., 2015. Is dispersal always beneficial to carrying capacity? new insights from the multi-patch logistic equation. Theoretical population biology 106, 45–59.

Arditi, R., Lobry, C., Sari, T., 2018. Asymmetric dispersal in the multi-patch logistic equation. Theoretical population biology 120, 11–15.

Belik, V., Geisel, T., Brockmann, D., 2011. Natural human mobility patterns and spatial spread of infectious diseases. Physical Review X 1 (1), 011001.

Bullo, F., 2024. Lectures on Network Systems, version 1.7 Edition. CreateSpace, http://motion.me.ucsb.edu/book-lns.

Butler, G., Freedman, H. I., Waltman, P., 1986. Uniformly persistent systems. Proceedings of the American Mathematical Society 96 (3), 425–430.

Cantrell, R. S., Cosner, C., 2004. Spatial Ecology via Reaction-Diffusion Equations. Wiley.

Capasso, V., Maddalena, L., 1981. Convergence to equilibrium states for a reaction-diffusion system modelling the spatial spread of a class of bacterial and viral diseases. Journal of Mathematical Biology 13 (2), 173–184.

Capitán, J. A., Cuenda, S., Alonso, D., 2017. Stochastic competitive exclusion leads to a cascade of species extinctions. Journal of Theoretical Biology 419, 137–151.

Chesson, P., 1994. Multispecies competition in variable environments. Theoretical population biology 45 (3), 227–276.

Colizza, V., Pastor-Satorras, R., Vespignani, A., 2007. Reaction–diffusion processes and metapopulation models in heterogeneous networks. Nature Physics 3 (4), 276–282.

Dieckmann, U., Ferrière, R., 2004. Adaptive Dynamics and Evolving Biodiversity. Cambridge Studies in Adaptive Dynamics. Cambridge University Press, p. 188–224.

Dockery, J., Hutson, V., Mischaikow, K., Pernarowski, M., 1998. The evolutionary stable strategy of dispersal and the ideal free distribution in a patchy environment. Journal of Mathematical Biology 37 (1), 61–83.

Erlander, S., Stewart, N. F., 1990. The gravity model in transportation analysis: theory and extensions. Vol. 3. Vsp.

Gjini, E., Madec, S., 2021. The ratio of single to co-colonization is key to complexity in interacting systems with multiple strains. Ecology and Evolution 11 (13), 8456–8474.

Grant, A. J., Restif, O., McKinley, T. J., Sheppard, M., Maskell, D. J., Mastroeni, P., 2008. Modelling within-host spatiotemporal dynamics of invasive bacterial disease. PLoS biology 6 (4), e74.

Gyllenberg, M., Hastings, A., Hanski, I., 1997. Structured metapopulation models. In: Metapopulation biology. Elsevier, pp. 93–122.

Halkola, A. S., Aittokallio, T., Parvinen, K., 2022. Tumor microenvironment as a metapopulation model: The effects of angiogenesis, emigration and treatment modalities. Journal of Theoretical Biology 545, 111147.

Hanski, I., 1991. Single-species metapopulation dynamics: concepts, models and observations. Biological journal of the Linnean Society 42 (1-2), 17–38.

Hanski, I., 1994. A practical model of metapopulation dynamics. Journal of Animal Ecology 63 (1), 151–162.

Hanski, I., 1999. Metapopulation ecology. Oxford University Press.

Hanski, I., Gilpin, M., 1991. Metapopulation dynamics: brief history and conceptual domain. Biological journal of the Linnean Society 42 (1-2), 3–16.

Hanski, I., Ovaskainen, O., 2000. The metapopulation capacity of a fragmented landscape. Nature 404 (6779), 755–758.

Hofbauer, J., Schreiber, S. J., 2022. Permanence via invasion graphs: Incorporating community assembly into modern coexistence theory. Journal of Mathematical Biology 85 (5), 54.

Holyoak, M., Leibold, M. A., Holt, R. D., 2005. Metacommunities: spatial dynamics and ecological communities. University of Chicago Press.

Keeling, M. J., Bjørnstad, O. N., Grenfell, B. T., 2004. Metapopulation dynamics of infectious diseases. In: Ecology, genetics and evolution of metapopulations. Elsevier, pp. 415–445.

Kraemer, M. U. G., et al., 2016. Progress and challenges in infectious disease cartography. Trends in Parasitology 32 (1), 19–29.

Le, T. M. T., Gjini, E., Madec, S., 2023. Quasi-neutral dynamics in a coinfection system with n strains and asymmetries along multiple traits. Journal of Mathematical Biology 87 (3), 48.

Le, T. M. T., Madec, S., 2023. Spatiotemporal evolution of coinfection dynamics: a reaction–diffusion model. Journal of Dynamics and Differential Equations, 1–46.

Leibold, M. A., Chase, J. M., 2018. Metacommunity ecology, volume 59. Princeton University Press.

Leibold, M. A., Holyoak, M., Mouquet, N., Amarasekare, P., Chase, J. M., Hoopes, M. F., Holt, R. D., Shurin, J. B., Law, R., Tilman, D., et al., 2004. The metacommunity concept: a framework for multi-scale community ecology. Ecology letters 7 (7), 601–613.

Lerch, B. A., Rudrapatna, A., Rabi, N., Wickman, J., Koffel, T., Klausmeier, C. A., 2023. Connecting local and regional scales with stochastic metacommunity models: Competition, ecological drift, and dispersal. Ecological Monographs 93 (4), e1591.

Levin, S. A., 1974. Dispersion and population interactions. The American Naturalist 108 (960), 207–228.

Levins, R., 1969. Some demographic and genetic consequences of environmental heterogeneity for biological control. Bulletin of the ESA 15 (3), 237–240.

Lin, H., Wen, T. H., 2022. How spatial epidemiology helps understand infectious disease transmission. Tropical Medicine and Infectious Disease 7 (8), 164.

Loreau, M., Mouquet, N., Holt, R. D., 2003. Meta-ecosystems: a theoretical framework for a spatial ecosystem ecology. Ecology Letters 6 (8), 673–679.

Madec, S., Gjini, E., 2020. Predicting n-strain coexistence from co-colonization interactions: epidemiology meets ecology and the replicator equation. Bulletin of Mathematical Biology 82 (142).

Madec, S., Gjini, E., 2025. Derivation of a spatial replicator system with environmental heterogeneity from a co-colonization sis model with n strains and p patches. submitted.

Malagon, D. A., Camper, B. T., Bewick, S., 2025. Multiscale metapopulations. Theoretical Ecology 18 (1), 32.

McCormack, R. K., Allen, L. J., 2007. Multi-patch deterministic and stochastic models for wildlife diseases. Journal of biological dynamics 1 (1), 63–85.

Meentemeyer, R. K., Haas, S. E., Vaclavik, T., 2012. Landscape epidemiology of emerging infectious diseases in natural and human-altered ecosystems. Annual Review of Phytopathology 50, 379–402.

Murray, J. D., Stanley, E. A., Brown, D. L., 1986. On the spatial spread of rabies among foxes. Proceedings of the Royal society of London. Series B. Biological sciences 229 (1255), 111–150.

Neuhauser, C., Pacala, S. W., 1999. An explicitly spatial version of the lotka-volterra model with interspecific competition. The Annals of Applied Probability 9 (4), 1226–1259.

Noble, J. V., 1974. Geographic and temporal development of plagues. Nature 250 (5469), 726–729.

Okubo, A., Levin, S. A., et al., 2001. Diffusion and ecological problems: modern perspectives. Vol. 14. Springer.

Ovaskainen, O., Rybicki, J., Abrego, N., 2019. What can observational data reveal about metacommunity processes? Ecography 42 (11), 1877–1886.

Parratt, S. R., Numminen, E., Laine, A.-L., 2016. Infectious disease dynamics in heterogeneous landscapes. Annual Review of Ecology, Evolution, and Systematics 47, 283–306.

Plümper, T., Neumayer, E., 2023. Tobler’s law and wavefront patterns in the spatial spread of covid-19 across europe during the delta and omicron waves. Scandinavian Journal of Public Health 51 (5), 775–779.

Poggiale, J.-C., Auger, P., Nerini, D., Manté, C., Gilbert, F., 2005. Global production increased by spatial heterogeneity in a population dynamics model. Acta Biotheoretica 53 (4), 359–370.

Pullan, R. L., Sturrock, H. J. W., Soares Magalhaes, R. J., Clements, A. C. A., Brooker, S. J., 2012. Spatial parasite ecology and epidemiology: a review of methods and applications. Parasitology 139 (14), 1870–1887.

Real, L. A., Biek, R., 2007. Spatial dynamics and genetics of infectious diseases on heterogeneous landscapes. Journal of the Royal Society Interface 4 (16), 935–948.

Skellam, J. G., 1951. Random dispersal in theoretical populations. Biometrika 38 (1/2), 196–218.

Smith, D. L., Dushoff, J., Perencevich, E. N., Harris, A. D., Levin, S. A., 2004. Persistent colonization and the spread of antibiotic resistance in nosocomial pathogens: resistance is a regional problem. Proceedings of the National Academy of Sciences 101 (10), 3709–3714.

Smith, H., 1995. Monotone Dynamical Systems: An Introduction to the Theory of Competitive and Cooperative Systems: An Introduction to the Theory of Competitive and Cooperative Systems. Mathematical surveys and monographs. American Mathematical Society. URL https://books.google.pt/books?id=vOfNAwAAQBAJ

Smith, H., Waltman, P., 1995. The Theory of the Chemostat: Dynamics of Microbial Competition. Cambridge Studies in Mathematical Biology. Cambridge University Press. URL https://books.google.pt/books?id=wFLdVo89vq8C

Steiner, M. C., Novembre, J., 2022. Population genetic models for the spatial spread of adaptive variants: A review in light of sars-cov-2 evolution. PLOS Genetics 18 (9), e1010391.

Tao, Y., Hastings, A., Lafferty, K. D., Hanski, I., Ovaskainen, O., 2024. Landscape fragmentation overturns classical metapopulation thinking. Proceedings of the National Academy of Sciences 121 (20), e2303846121.

Thompson, P. L., Guzman, L. M., De Meester, L., Horváth, Z., Ptacnik, R., Vanschoenwinkel, B., Viana, D. S., Chase, J. M., 2020. A process-based metacommunity framework linking local and regional scale community ecology. Ecology letters 23 (9), 1314–1329.

van Baalen, M., Sabelis, M. W., 1995. The dynamics of multiple infection and the evolution of virulence. The American Naturalist 146 (6), 881–910.

Wong, J. K., Yukl, S. A., 2016. Tissue reservoirs of hiv. Current Opinion in HIV and AIDS 11 (4), 362.

Zilio, G., Duncan, A. B., Ezenwa, V. O., Fenton, A., Gjini, E., Kaltz, O., Magalhães, S., Michalakis, Y., Mideo, N., Pedersen, A., et al., 2026. Co-circulation and co-infection: parasite interactions across scales. Trends in Ecology & Evolution.

Zipf, G. K., 1946. The p1 p2/d hypothesis: On the intercity movement of persons. American Sociological Review 11 (6), 677–686.

